# Experimental and simulated FRAP for the quantitative determination of protein diffusion in helical cells

**DOI:** 10.64898/2026.02.27.708671

**Authors:** Shariful Sakib, Cécile Fradin

## Abstract

Fluorescence recovery after photobleaching (FRAP) is widely used to characterize diffusion in cells, but quantitative interpretation of the data in small prokaryotes requires explicitly accounting for cell geometry. While this has been successfully achieved for spherical and rod-shaped bacteria, analytical approaches developed in these cases are not directly applicable to cells with more complex morphologies. Here, we explore the application of FRAP to helical bacteria using simulations. We show that half-compartment FRAP experiments, where one-half of the cell is photobleached, provide a robust means of characterizing fast protein diffusion. To help with the practical implementation of this technique, we established the relationship between the diffusion coefficient and characteristic fluorescence recovery time as a function of cell length and helical parameters, and for two different ways of estimating the recovery time. As a first application, we report measurements of the diffusion coefficient of the fluorescent protein, mNeonGreen, in the helical bacterium *Paramagnetospirillum magneticum* AMB-1. We find it to be *D* = 4.9 ± 2.2 µm^2^ s^−1^ in isosmotic conditions, not significantly different from the value measured in *Escherichia coli*. Although developed for helical bacteria, including spirilla, spirochetes, and vibrios, our framework can readily be extended to cells or compartments with other geometries.

## 1 INTRODUCTION

The small size of bacteria makes the quantification of cytoplasmic protein diffusion challenging in these organisms (1). Soluble proteins often diffuse too rapidly to be captured by single particle tracking (2), and the small dimensions of the bacterial cytoplasm, comparable to the diffraction limit of visible light, make fluorescence correlation spectroscopy difficult to implement (3). This leaves fluorescence recovery after photobleaching (FRAP) as the most viable method for studying cytoplasmic diffusion in bacteria. However, when applied to such small compartments, where the photobleached region can represent a substantial fraction of the cell volume, quantitative FRAP analyses become strongly dependent on the three-dimensional geometry of the cell (4).

FRAP has enabled precise measurements of intracellular protein diffusion in bacteria with simple morphologies, such as spherical cocci and cylindrical bacilli, most notably in *Escherichia coli* (*E. coli*) (3, 5–9). It has also been used to quantitatively probe not just diffusion but also molecular interactions and exchanges in other types of spherical compartments, for example protein-DNA binding and nucleocytoplasmic transport in cell nuclei (10–12), and rates of exchange into and out of biomolecular condensates (13–15). In contrast, in helical cells, FRAP has so far been limited to qualitative or comparative studies of bound or slow-diffusing proteins, in which case the helical geometry is irrelevant. For example, in magnetotactic spirilla, FRAP has helped establish the dynamics of the filament formed by the actin-like protein MamK (16–18), as well as those of the associated proteins MamJ, McaA, and McaB (18, 19). However, to our knowledge, FRAP has not yet been used to measure the diffusion coefficient of fast, freely-diffusing proteins in helical bacteria.

A common experimental strategy for FRAP experiments in confined volumes is to perform half-compartment bleach experiments, where roughly half of the compartment is targeted during the photobleaching step (5, 7, 14, 20). The resulting data can be analyzed either by considering the recovery of the fluorescence signal integrated over the bleached region (hereafter referred to as bleach region analysis)(13, 14, 20, 21), or by following the temporal decay of the amplitude of the first Fourier mode of the fluorescence intensity profile (hereafter referred to as Fourier mode analysis)(5, 7, 21, 22). For a single population of diffusing fluorophores, the fluorescence profile obeys the diffusion equation and the amplitudes of Fourier modes decay exponentially (5, 23). When the fluorescence profile is dominated by a single mode, the bleach region intensity should therefore also display an approximately exponential recovery. This was shown to be the case for impermeable spherical compartments, where fluorescence profiles can be decomposed into spherical Bessel functions (14, 24), and the overall fluorescence recovery in the bleached half-compartment is well described by a single exponential (20).

An exponential process is characterized by a single quantity, the characteristic recovery time τ. Dimensional analysis predicts that τ scales as *R*^2^/*D*, where *D* is the diffusion coefficient of the fluorophores and *R* is a characteristic linear dimension related to the size of the bleached region, the size of the compartment, or both. For half-compartment FRAP experiments, the length of the bleach region is tied to the compartment caliper length *L*_*T*_. For this reason, we chose to define a dimensionless recovery time, 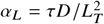, which removes the trivial dependence of τ on diffusion coefficient and compartment size, so that *α*_*L*_ should depend primarily on the geometry of the compartment and the bleach region. For simple geometries, *α*_*L*_ can be estimated analytically, by considering the recovery of the dominant mode of the fluorescence profile. For example, for a long cylindrical compartment, *α*_*L*_ = 1/*π*^2^ ≃ 0.101 for the first Fourier mode (5, 7), while for a sphere, *α*_*L*_ ≃ 0.0577 for the dominant spherical Bessel function (20). However, these analytical results do not readily extend to more complex geometries, such as the third most common bacterial shape, helical (spirillum-shaped) cells.

In this study, we used the well-characterized spirillum-shaped bacterium *Paramagnetospirillum magneticum*, formerly known as *Magnetospirillum magneticum* (hereafter referred to as AMB-1, for Aerobic Magnetotactic Bacteria) as a model system to establish a FRAP-based framework for studying protein mobility in cells with helical morphologies. We first present simulations of FRAP experiments in helical compartments designed to explore the relationship between the dimensionless recovery time that is represented by the coefficient, *α*_*L*_, and cellular geometry (helical pitch, helical amplitude, diameter and length), for both bleach region and Fourier mode analyses, and for different bleach strategies. Based on these simulations, we provide an empirical closed-form expression for *α*_*L*_ specific to half-compartment FRAP experiments, which offers a straightforward way to estimate the fluorophore diffusion coefficient, 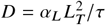. As a first application of this framework, we compare the diffusion coefficient of the monomeric yellow-green fluorescent protein, mNeonGreen (mNG), in AMB-1 and in *E. coli*.

## 2 METHODS

### 2.1 FRAP simulations

A computationally-efficient program that leverages the NumPy library in Python was written to simulate the diffusion of point particles within any volume that can be defined as the ensemble of points within a distance *r* from a user-defined curve. The curve is discretized into an array of points spaced by a distance *δs*, allowing for quick estimates of the minimum distance between any given point and the curve. For all simulations presented here, the point spacing was chosen as 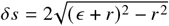, to limit the distance error to *ϵ* = 4.2 nm (roughly the length of a typical *β*-barreled fluorescent protein). We used this framework to simulate diffusion in helical cells defined by a discretized helical curve (length *L*, amplitude *A*, pitch *λ*). Setting *A* = 0 or *λ* = *∞* produces cylindrical compartments with two end caps of radius *r* that simulate the cytoplasmic geometry of rod-shaped cells, while setting *A* > 0 and *λ* < *∞* produces helical compartments that simulate the cytoplasmic geometry of spirillum-shaped cells (see schematics in Fig. 1).

**Figure 1.**
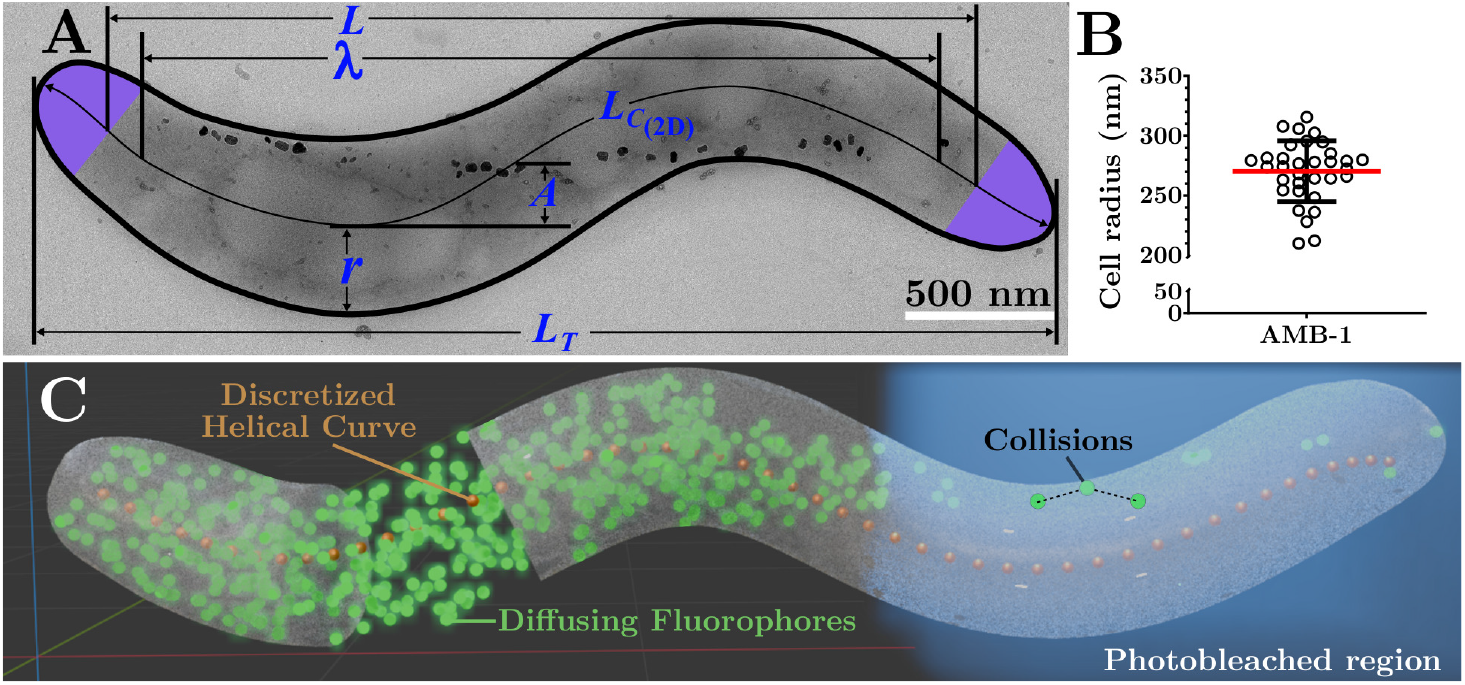
Helical compartment geometry. (A) Transmission electron micrograph of an AMB-1 cell annotated with the helical parameters used and defined in this study. B) AMB-1 cell radii, *r*, determined from TEM images. The average value *r* = 0.27 µm was used in the simulations and for experimental data analysis. C) 3D model of an AMB-1 cell created using Blender 4.4 illustrating the key components of the described simulated FRAP experiments.

At the beginning of the simulation, *N* = 1000 particles are randomly placed in the compartment, then allowed to diffuse within its confines as follows. At each time step *δt*, each particle is assigned a potential displacement vector 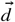 with components drawn from a zero-mean normal distribution with variance *σ*^2^ = 2*Dδt*. The distance between the potential new position of the particle and the helical curve is approximated as 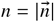 where 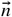 is the vector pointing from the particle to the closest point on the discretized curve. A collision event is triggered if this distance exceeds the value of the compartmental radius (*n* > *r*). In that case, 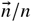 represents the unit vector normal to the cell membrane at the point of collision to a very good approximation as long as 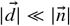. A corrected displacement vector is then calculated as 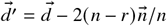 to simulate particle reflection off the cell membrane. For all simulations presented here, *δt* = 1 ms.

After a set time interval (150 ms), a photobleaching step is implemented for a short duration Δ*t*_bleach_ (30 ms), in a defined bleach region. In the simulations presented here, the bleach region was a cuboid of length *R*_*ℓ*_. For long cells (*L*_*T*_ > *λ*/2), it was oriented along the helical axis, whereas for short cells (*L*_*T*_ < *λ*/2) it was oriented along the line connecting the two end points of the discretized helical curve. This choice ensures that the bleach region is aligned with the apparent long axis of the cell, as it would be defined by a user from an image of the cell. At each time increment during the photobleaching step, each particle found in the bleach region is given a probability *p*_bleach_ = *δt*/*λ*_bleach_ of being deleted. *λ*_bleach_ is the fluorophore lifetime under photobleaching conditions. Unless otherwise specified, in the simulations presented here *γ* = Δ*t*_bleach_ /*λ*_bleach_ = 5. This value is high enough to ensure that the survival probability of a fluorophore remaining in the bleach region for Δ*t*_bleach_, given by *e*^−*γ*^, is low (strong photobleaching). It also ensures that *p*_bleach_ = *γδt*/Δ*t*_bleach_ ≪ 1, a necessary condition for correctly simulating progressive fluorophore photobleaching.

To reflect the sampling rate of the accompanying FRAP experiments, the simulation data was sampled at regular time intervals, Δ*t*_frame_ = 30 ms. The simulation was run for a time 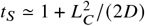 to ensure that the full fluorescence recovery was captured. The fluorescence in the photobleached and non-photobleached compartments was recorded as the count of particles present in each of these regions. The fluorescence profile along the cell axis was obtained by binning the particle count into 85 nm wide regions. The simulation data was then analyzed exactly in the same way as the experimental data to extract the fluorescence recovery time (τ), as detailed in section 2.5.

A video demonstrating the simulation of a FRAP experiment can be found in the Supplementary Material.

### 2.2 Cell culture

AMB-1 cells (purchased from the American Type Culture Collection, ATCC 700264) were grown in the magnetic spirillum growth medium described for *Magnetospirillum gryphiswaldense* strain MSR-1 by Le Nagard *et al*. (25). That medium, hereafter referred to as MSGM-LN (for Magnetospirillum Growth Medium - Le Nagard), was found to be optimal for the growth of AMB-1 when compared to other media described in the literature (see Supplementary Material for growth curves) (25–27). MSGM-LN was prepared by adding the following ingredients per litre of ddH_2_O: ATCC’s trace mineral supplement (product code: MD-TMS) 1 mL, ferric citrate (10 mM autoclaved stock) 5 mL, KH_2_PO_4_ 0.1 g, MgSO_4_ · 7H_2_O 0.15 g, HEPES 2.38 g, NaNO_3_ 0.34 g, yeast extract 0.1 g, soybean peptone (BD Bacto Soytone) 3.0 g, and potassium lactate (60% w/v) 4.35 mL. The pH was adjusted to 7. The medium was aliquoted into airtight serum bottles sealed with a rubber septum stopper and crimped with a metal cap, then sparged with N_2_ for 1 min/mL of media and autoclaved. AMB-1 cells were thawed from glycerol stock into MSGM-LN with a final 5 : 3 volumetric ratio of headspace to media, with the headspace being supplemented with O_2_ (first autoclaved in a separate serum bottle) with a syringe to a final concentration of approximately 1.5%. The initial lag-phase immediately after thawing typically lasted for 5 days until log phase began. At that time point, cells were subcultured via 1 : 20 dilution into a new serum bottle with fresh MSGM-LN media but with a 1 : 4 headspace-to-media ratio with atmospheric air (~21% O_2_) in the headspace (28). This growth regime balanced the O_2_ requirements for healthy growth of microaerophilic AMB-1 while also allowing the chromophore to mature within those cells. Cells were cultured for no more than a total of 10 days at a time from glycerol stock to ensure reproducibility.

*E. coli* strains TOP10 (Invitrogen) and MFDpir (kindly provided by Dr. Ghigo (29)) were used for cloning and conjugation, respectively. *E. coli* TOP10 cells were cultured in LB Miller broth while *E. coli* MFDpir cells were cultured in LB Lennox broth supplemented with 2,6-Diaminopimelic acid (DAP) to a final concentration of 0.3 mM. Standard incubation conditions of 37 °C and orbital shaking (250 RPM) were used with vented cell culture tubes with a headspace-to-media ratio of at least 3 : 1.

### 2.3 mNeonGreen expression in AMB-1

The low copy broad host range vector pRK415 (30, 31) with f-like plasmid features (*OriT* and *traJ*), and an *egfp* sequence under the control of a *lac* promoter was obtained from Nova Lifetech Inc. This genetic system allowed for low levels of protein expression to minimize the metabolic strain of non-native expression in AMB-1. A constitutive expression vector, pRK415-mNG, was created by replacing the *egfp* gene with the *mNG* gene. A vector backbone fragment was obtained by double digestion of pRK415-EGFP with 10 units each of EcoRI and BamHI in the rCutSmart™ buffer to excise the *egfp* sequence. A gBlocks™ Gene Fragment was chemically synthesized (Integrated DNA Technologies) to obtain the *mNG* sequence with homologous arms compatible with isothermal DNA assembly with the pRK415 backbone. The two fragments were then assembled using the GeneArt™ Gibson Assembly HiFi 2X Master Mix in a standard 50 °C isothermal reaction. An inducible variation of this vector with mNG expression under the control of a *tac* promoter and LacI repressor system was constructed via a more complex 5-fragment assembly that was optimized with a proprietary technique described in the Supplementary Material. The resultant pRK415-mNG constructs were used to transform chemically competent *E. coli* TOP10 cells (Thermo Fisher Scientific) or DH10Beta cells (New England Biolabs) via a standard heat shock procedure. The *E. coli* strains harbouring the plasmids are available through Addgene (See Data Availability Section). Plasmid DNA sequences were verified via Sanger sequencing.

For expression in AMB-1, the pRK415-mNG construct was shuffled into *E. coli* MFDpir cells that were made chemically competent by modifying the standard protocol outlined by Sambrook and Russell (32, 33), by including 0.3 mM DAP in both CaCl_2_ buffers. pRK415-mNG was then shuffled into AMB-1 via a standard conjugal transfer protocol for *magnetospirillum* (34) with a donor:recipient ratio of 1 : 1 and spot-plated onto EMSGM (35) 1.5% agar plates supplemented with antibiotic and DAP. The plate was kept sealed in a microaerophilic environment by using a BD GasPak™ EZ CampyPouch system with a water-moistened cotton ball and incubated at 30 °C for 6 h, after which, the conjugate mixture was plated once more on EMSGM 1.5% agar without DAP to select against the auxotrophic MFDpir strain. Colonies were picked and transferred to a serum bottle with fresh MSGM-LN culture media for about 4 days until log phase was reached. The culture media was then passed through a capillary racetrack filtration system (36) to select for healthy cells displaying the motility and magnetotaxis that is characteristic of this species.

When needed, the following antibiotics were used: Tetracycline (10 µg mL^−1^, 5 µg mL^−1^ for AMB-1), Streptomycin (100 µg mL^−1^), and Zeocin (50 µg mL^−1^).

### 2.4 FRAP experiments

#### 2.4.1 Sample preparation

For FRAP experiments *E. coli* and AMB-1 cells harbouring the pRK415-mNG plasmids were harvested during their respective log phases. AMB-1 cells subcultured in MSGM-LN with a 1 : 4 headspace-to-medium ratio as explained in section 2.2. At this time point immediately post subculturing (lag phase), the AMB-1 with pRK415-mNG (inducible expression) was induced with a final concentration of 0.1, 0.5, or 1 mM of IPTG to create a pseudo-constitutive expression regime to mimic the expression conferred by the constitutive expression vector. This allowed us to assay the effects of varying the strength of constitutive expression within AMB-1. All AMB-1 cells from each expression regime were harvested after 16 h. *E. coli* TOP10 (K-12 derivative) cells were first streaked and incubated on LB Miller 1.5% agar plates for 16 h. Colonies were picked and grown in LB Miller broth for another 12−16 h until an OD_600_ of 0.5 was attained. Cells were then subcultured in Terrific Broth and harvested after 3 h. Harvested cells (1.5 mL cell culture medium) were centrifuged at 1000×g for 5 min, and washed three times with 500 µL of Dulbecco’s Phosphate Buffered Saline (DPBS) with no MgCl_2_ or CaCl_2_, and with an osmolarity adjusted to that of the cells. For *E. coli* 1×DPBS (osmolarity of approximately 300 mOsm L^−1^) was used, while for AMB-1 DPBS was diluted to an osmolarity of 85 mOsm L^−1^ (see Supplementary Materials for detailed osmolarity calculations). The cells were then stained with Cellbrite® Membrane Fix 640 stain (1000× stock) at a final concentration of 10× at room temperature for 15 min while rocked at 30 RPM. Cells were centrifuged and washed in 500 µL of isosmotic DPBS three times to remove excess stain, then resuspended in isosmotic DPBS. An 8 µL volume of the cell suspension was pipetted onto a microscope slide and covered with an 18 × 18 mm No. 1.5 coverslip. All glassware was first rinsed with anhydrous ethanol and dried, before being coated with 0.01% Poly-L-Lysine (PLL) solution (facilitating gentle cell immobilization), then rinsed with DI water and dried again. Samples were quickly sealed with clear nail polish to avoid evaporation and imaged within 30 min to maintain high cell viability. Detailed visual instructions of the sample preparation procedure are provided in the Supplementary Material.

#### 2.4.2 Photobleaching and confocal imaging protocol

FRAP experiments were performed on a ZEISS LSM980 Inverted confocal microscope with the ZEN 3.8 software, using a 63X oil immersion objective (NA = 1.4) in conjunction with a 2 Airy unit pinhole. The fluorescence of mNG and CellBrite® Fix 640 was excited at 488 nm and 639 nm, respectively. Their fluorescence emissions were separated using wavelength range presets for mNG and AlexaFluor 647, and collected with GaAsP detectors (gain 800 V for both channels). For FRAP experiments, images with dimensions 6.73 × 6.73 µm^2^ (100 × 100 px^2^) were acquired in the mNG channel, with a pixel dwell time of 0.6256 µs, a frame generation time of 14.64 ms, and a frame interval of approximately 30 ms. A rectangular bleach region was defined with the ZEN software to cover approximately half of the cell. Photobleaching was performed over a single frame interval using 50% of the maximal laser power (5 out of 10 mW for the 488 nm and 6.5 out of 13 mW for the 639 nm laser lines), preceded and followed by imaging at 0.35% of the maximal power. The total duration of a FRAP experiment was set to 3 seconds for each bacterium. All FRAP experiments were conducted at room temperature, *T* = 23 ± 2 °C.

#### 2.4.3 Automated image processing

Image analysis for FRAP experiments was automated using PyImageJ, a Python interface that initializes a Java Virtual Machine to control ImageJ and execute macros (37) (as illustrated in Fig. 2). The manually defined bleach region is imported using the bioformats ImageJ plugin, and other relevant parameters (frame interval, photobleach time index, duration of photobleaching step) are extracted from the metadata via scripting. Several regions of interest (ROIs) are automatically defined. ROI I is the rectangular user-defined bleach region. ROI II comprises all pixels within the cell body. It is specified by averaging the images recorded prior to the photobleaching step, filtering the average image with a Gaussian blur (*σ* = 3 pixels) then thresholding the filtered image. The preset Yen thresholding algorithm is used by default, though in the presented experiments a small number of AMB-1 cells (6 out of 67) required the Triangle algorithm for correct thresholding. Cell caliper length, *L*_*T*_, is measured by using the ImageJ Rectangle Fit function to draw a bounding box around the cell (ROI III). Two masks corresponding to the bleached (ROI IV) and non-bleached (ROI V) regions of the cell are created with Fiji’s logical ROI operator functions: ROI IV is obtained from ROIs I and II using the “and” (&&) operator, while ROI V is obtained from ROIs IV and II by using the “exclusive or” (XOR) operator. The average fluorescence over time within the bleached region, *I*_*b*_(*t*), and non-bleached region, *I*_*n*_(*t*), are computed using the ImageJ Plot Z-axis Profile function. The fluorescence intensity profile along the cell’s long axis at each time point (*i*(*x, t*)) is automatically calculated as the average pixel intensity for each column of pixels in ROI III using the ImageJ Plot Profile function. These intensities are exported in CSV format for automated data analysis.

**Figure 2.**
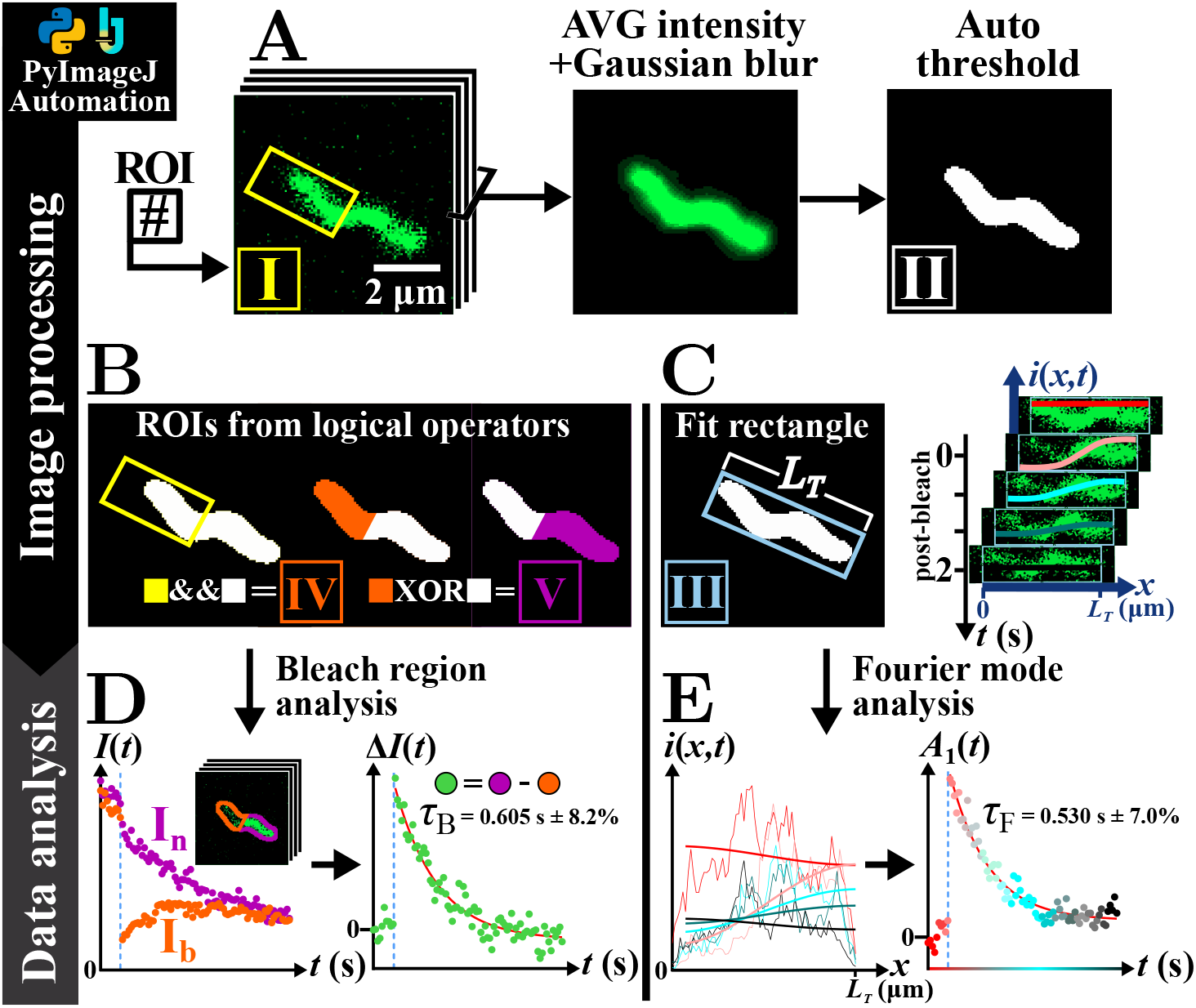
FRAP image processing and data analysis workflow. Example of the PyImageJ processing and analysis of a representative half-compartment FRAP experiment performed on an AMB-1 cell expressing mNG. (A) The image stack and position of the bleach region (ROI I) are imported. The average pre-bleach image is filtered and thresholded to define the region corresponding to the cell body (ROI II). (B) The cell’s bleached and non-bleached regions (ROIs IV and V) are defined through logical operations. (C) The cell body bounding rectangle (ROI III) is used to calculate the total cell length, *L*_*T*_, and to generate a fluorescence intensity profile at each time point. (D) Bleach region analysis is performed by computing the total fluorescence intensity in ROIs IV and V, then fitting the difference, Δ*I*(*t*), to an exponential function. (E) Fourier mode analysis is performed by fitting the intensity profiles to a cosine function, and then the amplitude of this cosine function, *A*_1_ (*t*), to an exponential decay. (D,E) Both analysis methods return a value for the characteristic recovery time, τ.

### 2.5 Extraction of characteristic recovery times after photobleaching

Both the experimental and the simulated FRAP data were analyzed using Python scripts to extract characteristic fluorescence recovery times. For each data set, two separate methods were applied, yielding two different recovery times.

#### 2.5.1 Bleach region analysis

The difference in intensity between the non-bleached and bleached regions, Δ*I*(*t*) = *I*_*n*_ (*t*) − *I*_*b*_(*t*) was calculated and fit to the exponential function:

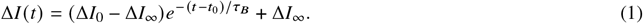

The fit is performed for *t* ≥ *t*_0_, where *t*_0_ is the first time point recorded immediately after the photobleaching step for both the simulations and experiments. It returns a value for τ_*B*_, which is the characteristic recovery time associated with the fluorescence integrated over the bleach region. It also returns the initial intensity difference between bleached and non-bleached regions, Δ*I*_0_, and any residual long-term intensity discrepancy between these two regions, Δ*I*_*∞*_.

#### 2.5.2 Fourier mode analysis

For each time point after photobleaching, the fluorescence intensity profile along the cell axis, excluding the semi-spherical caps at each end and thus spanning only a length *L* = *L*_*T*_ − 2*r*, was fitted to the cosine function: *i*(*x, t*) = *A*_0_(*t*) + *A*_1_(*t*) cos(*πx*/*L*_*T*_). This yields the amplitude of the first Fourier mode of the intensity profile, *A*_1_(*t*), which is then fitted to an exponential function:

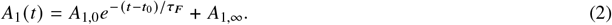

This returns a value for τ_*F*_, the characteristic relaxation time of the first Fourier mode.

### 2.6 Transmission Electron Microscopy

In preparation for Transmission Electron Microscopy (TEM) experiments, AMB-1 cells were harvested during log phase, washed three times with isotonic DPBS, and fixed with 0.01% formaldehyde for 10 min at 4 °C. Next, 4 µL of the fixed cells were transferred to a 400 mesh Au TEM grid coated with a carbon film, allowed to sit for 10 min, and washed with a stream of isotonic DPBS. The grids were then air-dried for another 10 min. For each imaged bacterium, a series of (2048 × 2048 pixels, 0.8119 nm pixel size, 1 s exposure) images were acquired with a TALOS L120C instrument equipped with a 4K × 4K Thermo Scientific Ceta CMOS camera. These images were then stitched together using an ImageJ plugin created by Preibisch *et al*. (38). The internal radius of the cell was then determined using ImageJ.

## 3 RESULTS

### 3.1 Interpretation of FRAP experiments in helical compartments

#### 3.1.1 Simulation of FRAP experiments in helical cells

To establish a quantitative framework for the interpretation of FRAP experiments in helical bacteria, we simulated FRAP for fluorophores diffusing within helical compartments. A representative simulation is shown in Fig. 3, with in this case helical parameters chosen to match those of typical AMB-1 cells: helical pitch *λ* = 2.5 µm, helical amplitude *A* = 0.2 µm, internal radius *r* = 0.27 µm. Values for *λ* and *A* were previously obtained from the analysis of light microscopy images of AMB-1 cells (39), while the value of *r* was measured from TEM images (*r* = 0.27 ± 0.03 µm, mean ± standard deviation (SD), *n* = 33 cells, Fig. 1B) - these parameters are relatively constant in AMB-1 cells. The caliper length of the simulated cell, *L*_*T*_ = 4.6 µm (including two end caps of radius *r*) and fluorophore diffusion coefficient, *D* = 4.4 µm^2^ s^−1^, match what is observed for the bacterium shown in Fig. 2. The bleach region in this simulation covers exactly one half of the cell (half-compartment bleach experiment), a common experimental choice for FRAP experiments in small compartments.

**Figure 3.**
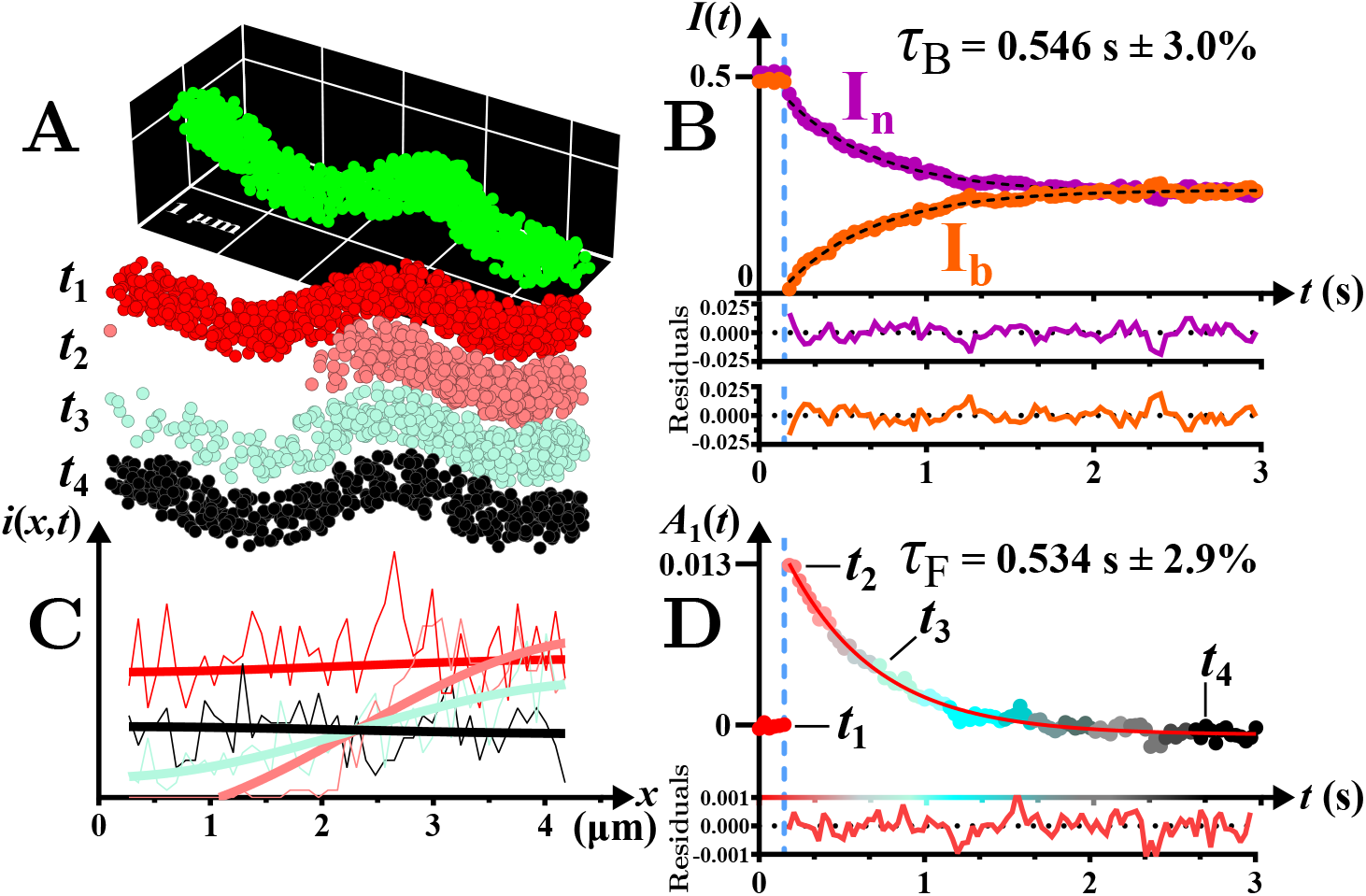
Half-compartment FRAP experiment simulation. Representative simulation for a helical cell with geometry mirroring that of the AMB-1 cell in Fig. 2. (A) Time points (*t*_1_ to *t*_4_) illustrate the cell pre-bleach state (red), immediately post-bleach (pink), and then the gradual transition towards fluorescence recovery (turquoise, black). (B) Result of bleach region analysis, with the fluorescence intensity in the bleached region (*I*_*b*_(*t*), orange symbols) and non-bleached region (*I*_*n*_(*t*), purple symbols) plotted against time. Black dashed lines are single-exponential fits, with characteristic decay time τ_*B*_. (C) Fluorescence intensity profiles, *i* (*x, t*), along the cell length, plotted for the time points shown in (A). Thick lines are fit to a cosine function (first Fourier mode). (D) Result of Fourier mode analysis, with the amplitude of the first Fourier mode, *A*_1_ (*t*), plotted as a function of time. The solid line is a fit to a single-exponential function, with characteristic decay time τ_*F*_. A video of the simulation can be found as a Supplementary Material.

Two methods were used to quantify fluorescence recovery: either monitoring the evolution of the fluorescence signal in the bleached and non-bleached regions (bleach region analysis, returning the characteristic time τ_*B*_, Fig. 2B), or alternatively considering the evolution of the amplitude of the first Fourier mode of the fluorescence profile (Fourier mode analysis, returning the characteristic time τ_*F*_, Fig. 2C,D). The simulated data (Fig. 3B,D) strongly resembles the experimental data (Fig. 2D,E), with the notable difference that photobleaching is observed during frame acquisition in FRAP experiments. Despite the complex helical geometry of the simulated compartment, both the fluorescence recovery observed for the bleach region (Fig. 3B), and the relaxation of the first Fourier mode over time (Fig. 3D), are undistinguishable from an exponential decay. This justifies fitting both real and simulated FRAP data of single diffusing-species in helical compartments with single exponential functions, and concentrating our discussion around the decay times τ_*B*_ and τ_*F*_ characterizing the fluorescence recovery process. In this example, the values of the characteristic recovery times obtained from bleach region and Fourier mode analysis are very similar.

#### 3.1.2 Influence of compartment aspect ratio on characteristic recovery time

Simulations allowed us to explore how fluorescence recovery is affected by cellular geometry, starting with changes associated with cell length. Within a bacterial population, body length naturally varies as each cell finds itself at a different stage of its growth cycle. Healthy *E. coli* cells for example have *L*_*T*_ ≈ 1 to 4.5 µm (40), and AMB-1 cells *L*_*T*_ ≈ 1.8 to 4.2 µm (39, 41). Meanwhile, the diameter of these cells is relatively constant (*d* ≃ 0.6 − 1 µm for *E. coli* (40, 42, 43), *d* = 0.5 − 0.6 µm for AMB-1). Thus, as bacteria grow, their aspect ratio changes and we expect a change in the value of the dimensionless recovery time 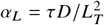.

To first explore how compartment aspect ratio influences fluorescence recovery in a simple case, we simulated half-compartment FRAP experiments for cylindrical cells of various internal diameters and lengths, and calculated 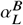 (or 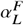) using the values of τ_*B*_ (or τ_*F*_) extracted from the simulations (Fig. 4A). To make the simulations more efficient, the particles’ diffusion coefficient was set to different values depending on compartment length: *D* = 1 µm^2^ s^−1^ for *L*_*T*_ ≤ 1 µm, 5 µm^2^ s^−1^ for 1 < *L*_*T*_ ≤ 5 µm and 15 µm^2^ s^−1^ for *L*_*T*_ ≥ 5 µm. As expected, we observe that, to first order, *α*_*L*_ depends on the geometric ratio *L*_*T*_ / *d* rather than on *D* and *L*_*T*_ independently. As *L*_*T*_ / *d* varies from 1 to 50, and as the compartment’s shape turns from a sphere to a long thin cylinder, passing through the pill-shaped morphology characteristic of *E. coli* when *L*_*T*_ / *d* ≃ 4, *α*_*L*_ monotonously increases. This can be understood as the increase in recovery time expected when diffusion goes from three-dimensional (spherical compartment, τ ~ ( *L*_*T*_ /2)^2^/(6*D*)) to one-dimensional (long cylindrical compartment, τ ~ ( *L*_*T*_ /2)^2^/(2*D*)).

**Figure 4.**
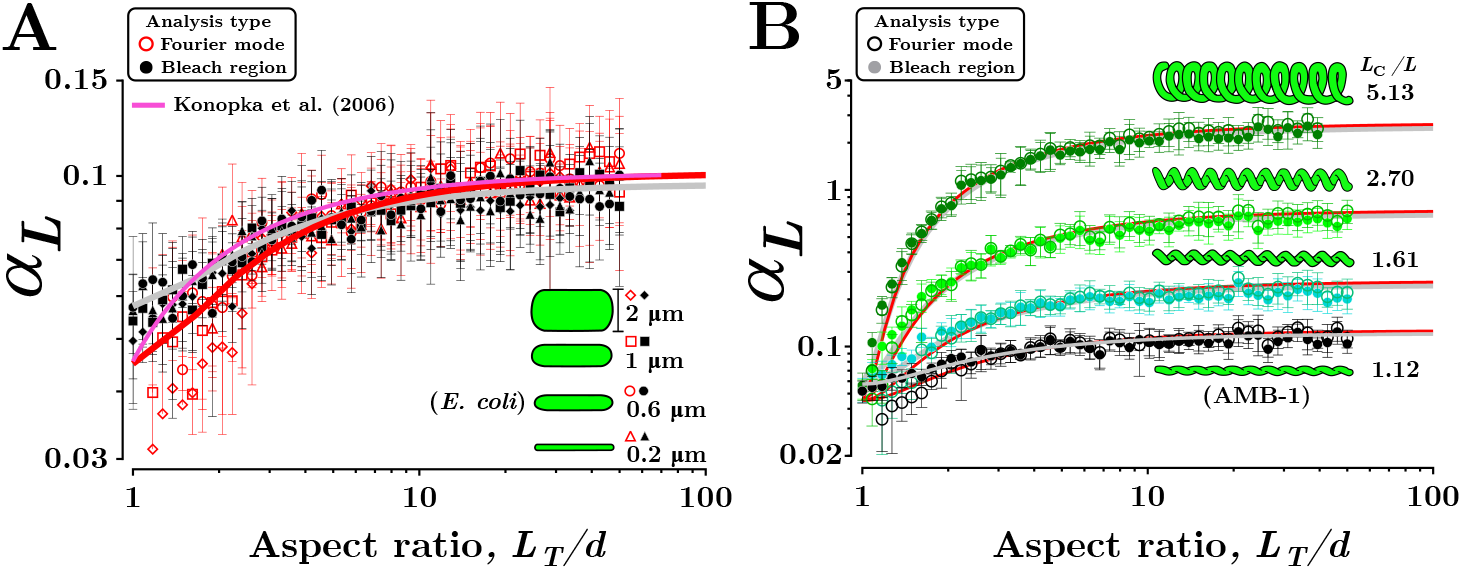
Influence of compartment aspect ratio and helicity on fluorescence recovery time. (A) Value of the dimensionless recovery time *α*_*L*_ obtained from simulated half-compartment FRAP experiments in cylindrical compartments with spherical caps as a function of compartment aspect ratio, *L*_*T*_ / *d*. triangle, circle, square, and diamond symbols represent values obtained for compartments with diameters *d* = 0.2, 0.6, 1, and 2 µm, respectively with Fourier mode data in red and bleach region data in black. The continuous pink line shows the approximate value of *α*_*L*_ proposed by Konopka *et al*. (7). (B) Value of *α*_*L*_ as a function of cell aspect ratio, *L*_*T*_ / *d*, obtained for different helical compartment geometries. Black, turquoise, light green and dark green symbols represent values obtained for compartments with helical amplitudes *A* = 0.2 µm (the value observed for AMB-1), 0.5, 1 and 2 µm respectively. The pitch (*λ* = 2.5 µm) and helical radius (*r* = 0.27 µm) were kept constant. In (A,B), filled symbols show values obtained using the bleach region analysis and empty symbols using the Fourier mode analysis. Grey and red lines show the form of the proposed phenomenological equation for 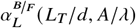 derived from all our simulations (Eq. 4) and act as visual guides for the simulated bleach region and Fourier mode analysis data, respectively. Each data point is the result of 10 replicate simulations (mean ± SD).

When *L*_*T*_ /*d* < 3, there is a noticeable difference between the values obtained for 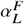 and 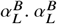 tends towards the value 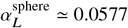 predicted for the dominant spherical Bessel function in the profile (20). In contrast 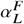 decreases towards a noticeably lower value. At the same time, the error on the estimated value of 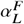 becomes very large. It is a sign that in this region reducing the fluorescence recovery to a one-dimensional profile analysis over a cylindrical portion that is becoming increasingly small in comparison to the compartment spherical end caps becomes unreliable. Although less noticeable, the two considered analysis methods also differ in the recovery time they return at large aspect ratios. While Fourier analysis returns values of 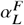 that exactly tends towards 1 / *π*^2^ (the value corresponding to the relaxation of the first Fourier mode (5)) at infinite *L*_*T*_ / *d*, the bleach region analysis returns slightly lower values for 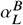. This is expected, since the fluorescence signal integrated over a sizable region becomes sensitive to a number of Fourier modes, which all have faster decays than the first Fourier mode. Fits of analytically calculated fluorescence decays for a cylinder predict that, when the fit of the exponential decay is carried out over a sufficiently large region as done in the simulations and experiments presented here, 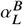 should tend towards 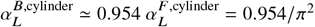 (see Supplementary material).

By considering a large range of aspect ratios (*L*_*T*_ /*d* = 1 to 50), and both common types of FRAP data analysis, our results refine and extend the results of a previous study in which the empirical formula 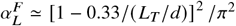 was proposed based on simulations performed in the range *L*_*T*_ /*d* = 1.8 to 3.5 relevant to *E. coli* cells (pink line in Fig. 4A) (7). Overall, our simulated data set highlight that it is very important to consider cell aspect ratio when calculating 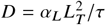 from the measured recovery time, and analysis method for cells with either very small or very large aspect ratios.

#### 3.1.3 Influence of compartment helicity on characteristic recovery time

The contour length of a helix of length *L* is given by *L*_*C*_ = *L* [1 + (2*π A* /*λ*) ^2^] ^1/2^. Thus recovery times (and in turn *α*_*L*_) should be larger for a helical cell than for a cylindrical cell of equal caliper length. In the limit of very long and thin compartments (quasi one-dimensional diffusion and negligible contribution from the end caps) we expect 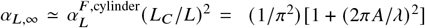. To confirm this intuition, we performed half-compartment FRAP experiments for helical cells with different helical amplitudes (starting with the average value for AMB-1 cells *A* = 0.2 µm). The values of *α*_*L*_ obtained from these simulations are shown in Fig. 4B. As is the case for cylindrical cells, *α*_*L*_ increases with compartment aspect ratio, from the value expected for a sphere at *L*_*T*_ / *d* = 1 to a plateau value when *L*_*T*_ / *d* ~ 10. This plateau value, *α*_*L, ∞*_, quickly increases as the helical amplitude, and thus the helix contour length, increases. As for the cylindrical cells, there are small systematic differences between 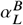 and 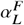, with 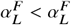 at short aspect ratios, and vice versa at large aspect ratios.

To test the dependence of *α*_*L, ∞*_ on *L*_*C*_ / *L*, we simulated FRAP experiments in helical compartments with a large fixed aspect ratio *L*_*T*_ / *d* = 30 (*L*_*T*_ = 16.2 µm, *d* = 0.54 µm) and extracted *α*_*L*, ∞_ while either varying the helical pitch (with fixed *A* = 0.2 µm, green symbols) or varying the helical amplitude (with fixed *λ* = 2.5 µm, black and red symbols). The results of these simulations, shown in Fig. 5A, confirm that *α*_*L, ∞*_ scales as (*L*_*C*_ / *L*) ^2^. A fit of the values obtained from the Fourier analysis (red line) show that they closely approach the expected (*L*_*C*_ / *L*) ^2^ / *π*^2^ (purple line), where the small 5% difference can be attributed to the fact that the aspect ratio used for the simulations, *L*_*T*_ / *d* = 30, is finite. A fit of the data obtained from bleach region analysis (grey line) show that these are on average about 10% smaller that those obtained from Fourier mode analysis, reflecting the capture of higher order and faster decaying Fourier modes mentioned above. Note that this linear relationship does not hold in the simulations when *λ* starts approaching the cell diameter, which for the series represented by green symbols in Fig. 5A happens for *L*_*C*_/*L* > 4.

**Figure 5.**
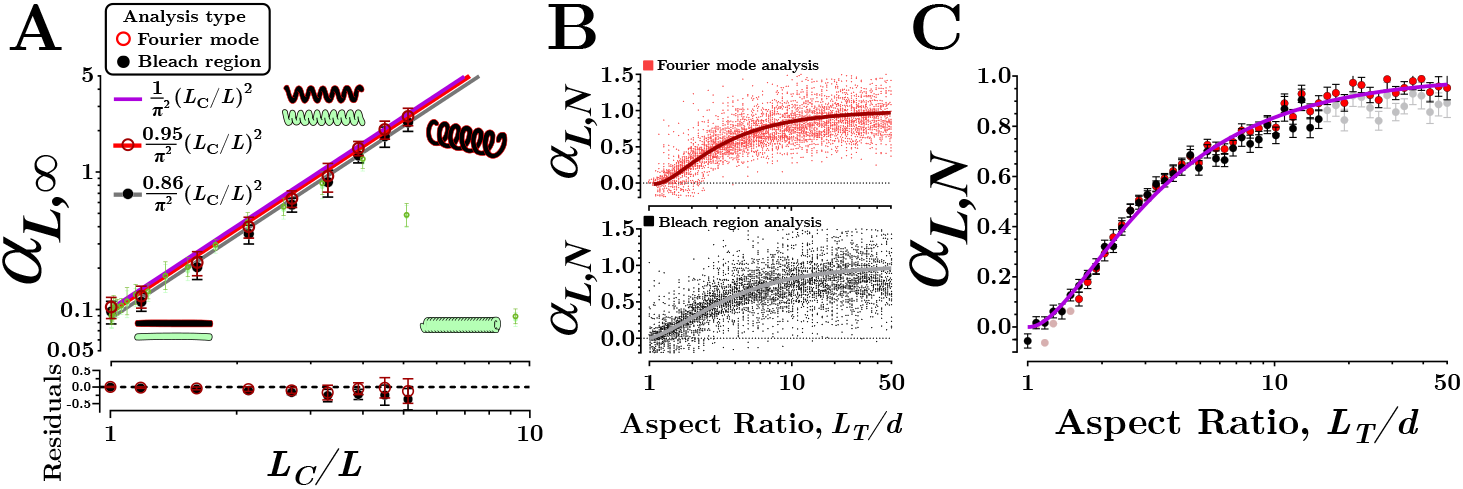
Universal form of *α*_*L*_ for both cylindrical and helical cells and both Fourier mode and bleach region analysis. (A) Value of *α*_*L*_ for helical cells with an aspect ratio *L*_*T*_/*d* = 30 as a function of normalized helix contour length, *L*_*C*_ / *L*. In the amplitude assay (black and red symbols), values for *A* were varied from 0 to 2 µm while holding the pitch constant (*λ* = 2.5 µm). In the pitch assay (pale green symbols), values for *λ* were varied from 0.68 to 100 µm while holding *A* constant (*A* = 1.0 µm). For all simulations *L*_*T*_ = 16.2 µm and *d* = 0.54 µm. The pink line represents the theoretical prediction *α*_*L,∞*_ = (1/*π*^2^)( *L*_*C*_/*L*)^2^. The dark blue and light blue lines represent fits of the Fourier mode and bleach region analysis data to the function (*C*/*π*^2^) × ( *L*_*C*_/*L*)^2^, returning *C* = 0.95 and 0.86, respectively. Each data point is the result of 10 replicate simulations (mean ± SD).(B) Normalized value of 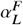 (top panel) and 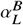 (top panel) denoted as 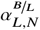, calculated as explained in the text from the simulation data in Fig. 4A,B obtained for both cylindrical and helical morphologies, as a function of compartment aspect ratio. The dispersion of the data is presented by showing all the replicates. Continuous lines represent the value of *α*_*L,N*_ that has been fitted with Eq. 3. (C) Averaged *α*_*L,N*_ as a function of compartment aspect ratio across morphologies for the Fourier mode analysis (red and light pink symbols) and the bleach region analysis (black and light grey symbols). Data points are mean ± SEM. The continuous purple line shows the fit of the averaged data to Eq. 3, taking into account both Fourier mode analysis and bleach region analysis data, but not the light pink and light grey data points that were considered less reliable.

#### 3.1.4 Universal relationship between diffusion coefficient and characteristic recovery time for cylindrical and helical cells

To obtain a universal relationship between *D* and τ, that would encompass cylindrical and helical cells, as well as Fourier mode and bleach region analysis, we considered the quantity *α*_*L,N*_ = [*α*_*L*_ − *α*_*L*,sphere_]/[*α*_*L,∞*_( *L*_*C*_/*L*)^2^ − *α*_*L*,sphere_], a normalized version of *α*_*L*_ calculated using values of *α*_*L*,sphere_ and *α*_*L,∞*_ adapted to each analysis type. For bleach analysis data, we used 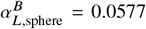 (first spherical Bessel function decay) and 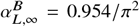 (predicted from integration of analytical intensity profiles). For Fourier mode analysis we used 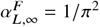 (first Fourier mode decay) and 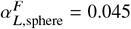 (obtained by minimizing the squared differences between 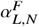 and 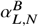). The normalized quantities 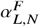 and 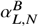 are first shown separately in Fig. 5B for all simulation repeats, giving an appreciation of the spread arising from both different cell geometries and experimental noise. The corresponding mean values are then compared in Fig. 5C. To a very good approximation, *α*_*L,N*_ values obtained for both analysis method collapse onto a single universal curve, well described by the analytical expression:

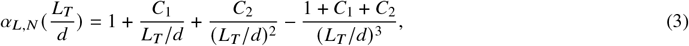

with *C*1 = −1.689, *C*2 = 0.357, and 1 + *C*1 + *C*2 = −0.332.

Thus the value of 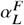 and 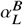 for cylindrical or helical cells of any dimensions can be approximated as:

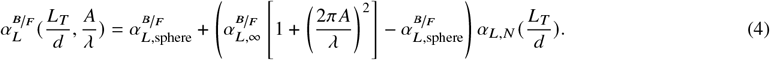

This equation provides a good prediction of the value of *α*_*L*_ for all the simulations we have performed. This can be seen in Fig. 4, where 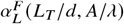 (red lines) and 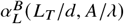 (grey lines) calculated using Eq. 4 have been plotted for all simulated datasets.

In practice, this means that if the caliper length (*L*_*T*_), body thickness (*d*), helical amplitude (*A*) and pitch (*λ*) of a helical cell body are known, the diffusion coefficient of a fluorophore can be estimated directly from the half-compartment bleach recovery time (τ) using the formula:

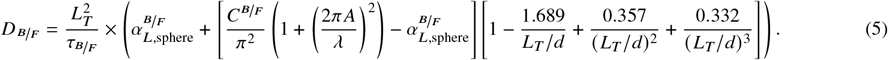

### 3.2 Guidelines for practical implementation of FRAP experiments in bacterial cells

In order to guide experimental choices, we then used simulations to explore the influence of the size and location of the bleach region along with the duration of the photobleaching step and frame acquisition interval.

#### 3.2.1 Influence of bleached region location and size

To determine the photobleaching geometry and analysis approach most conducive to robust FRAP measurements, we compared half-compartment FRAP experiments with central-region FRAP experiments, where a region at the cell center is bleached (Fig. 6). In the first case, the size of the terminal bleach region (*R*_*ℓ*_) was varied to simulate situations when the region defined by the user is not exactly equal to half the cell length (Fig. 6a). In the second case, both the position (*x*) and the size (*R*_*ℓ*_) of the bleached region were varied (Fig. 6b).

**Figure 6.**
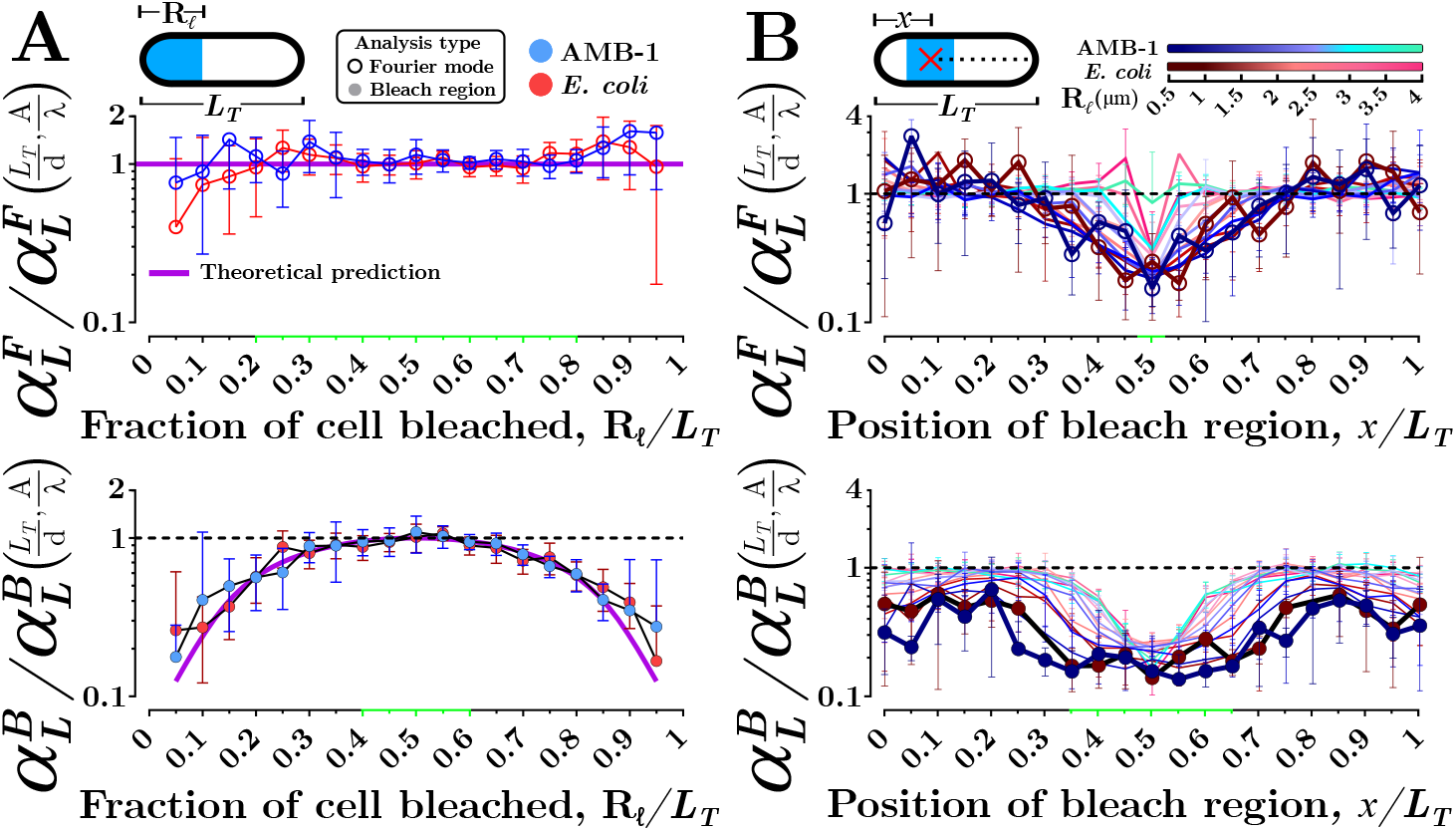
Dependence of recovery time on bleached region size and location. Normalized recovery time *α*_*L*_ as a function of (A) terminal bleach region size or (B) central bleach region location and size. Data was obtained from simulations of *L*_*T*_ = 5 µm compartments shaped either as a typical *E. coli* cell (cylindrical, *L*_*T*_ /*d* = 8.3 and *L*_*C*_/*L* = 1, red symbols and lines) or as a typical *P. magneticum* cell (helical, *L*_*T*_ /*d* = 9.3 and *L*_*C*_/*L* = 1.12, blue symbols and lines), using 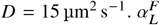 and 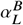 are separately shown in the top and bottom panels, respectively. All data was normalized with the value 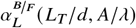 expected in the ideal half-compartment bleach case for the simulated cell and calculated with Eq. 4. In (A) purple lines represent the predicted value of *α*_*L*_ (see text for details). In (B) the length of the photobleached region, *R*_*ℓ*_, is reflected in symbols and lines color value (symbols are displayed only for the two smallest values of *R*_*ℓ*_). Green-shaded regions demarcate the conditions in which calculating *D* is robust against small variations in the length (A) or placement (B) of the bleach region. Each data point shows the mean ± SD of 10 simulation replicates.

Half-compartment bleach experiments are very robust against variation in the bleach region length, where the recovery time and the coefficient *α*_*L*_ used to calculate *D* varies very little as long as the terminally bleached region remains between 40 to 60% of the cell length (Fig. 6A). This holds true for both types of data analysis, but it is most striking when considering Fourier mode analysis, for which any bleached terminal region between 20 to 80% of the cell length results in the same normalized recovery time, 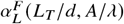 (represented by the purple line in the top panel of Fig. 6A) for a given cell geometry. This remarkable stability reflects the fact that the first Fourier mode captures the effect of perturbations on the scale of the entire cell, regardless of the exact length of the initial terminal perturbation (photobleach region). Bleach region analysis, on the other hand, involves integration of a signal that is sensitive to all Fourier modes, and as *R*_*ℓ*_ deviates from its ideal *L*_*T*_ / 2 value, higher order modes with faster decays become more prevalent, leading a marked decrease in 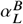. This decrease can be exactly predicted by integrating the analytically generated profiles (pink line in bottom panel of Fig. 6A, see Supplementary Materials for details).

The outlook is quite different when considering a central bleach region (Fig. 6B). In that case, both the position and size of the bleach region have a strong influence on the recovery time. For both Fourier mode and bleach region analyses, as the bleach region is moved from the edge of the compartment (a situation which is exactly equivalent to performing an imperfect half-bleach experiment) to the center of the compartment, no matter the value of *R*_*ℓ*_, *α*_*L*_ decreases by a factor of about 4. For 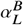 (bottom panel), this can be understood with a simple argument, which is that a small photobleaching area in the center of a cell with length *L*_*T*_ is equivalent to performing half-compartment bleach experiments in two adjacent cells of length *L*_*T*_ / 2 - leading to a 4-fold faster recovery (an effect discussed for example in Ref. (4)). Thus, small errors in the location of the photobleaching area in this type of experiment can lead to large errors in the determination of *D*. The first Fourier mode analysis (top panel), is not well adapted to central bleach region FRAP, because the amplitude of that mode is zero in the case of a perturbation located in the center of the cell. As a result the recovery times captured captured in this case using first Fourier mode analysis are very inconsistent. In conclusion, if performing FRAP using a central bleach region, one should ensure that the bounds of the central bleach region is within 40 to 60% of the cell’s center, that the region is as small as possible, and that bleach region analysis is performed.

#### 3.2.2 Influence of photobleaching step duration

FRAP experiments with prolonged photobleaching durations are susceptible to the so-called halo effect, in which the effective size of the bleached region increases as photobleached fluorophores diffuse out of the user-defined region while photobleaching is still in progress (44–46). If unaccounted for, this can lead to an incorrect estimate of *D*. In the quasi one-dimensional geometry of the bacterial cells considered here, the halo effect is expected to become significant when the photobleaching duration Δ*t*_bleach_ exceeds 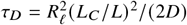, the timescale over which a particle can diffuse in our out of the bleach region. We tested for this prediction for both half-compartment and central region FRAP assays by extracting *α*_*L*_ from simulations in which Δ*t*_bleach_ was systematically varied within a range of values (0 to 500 ms) corresponding to what could typically be achieved experimentally (Fig. 7).

**Figure 7.**
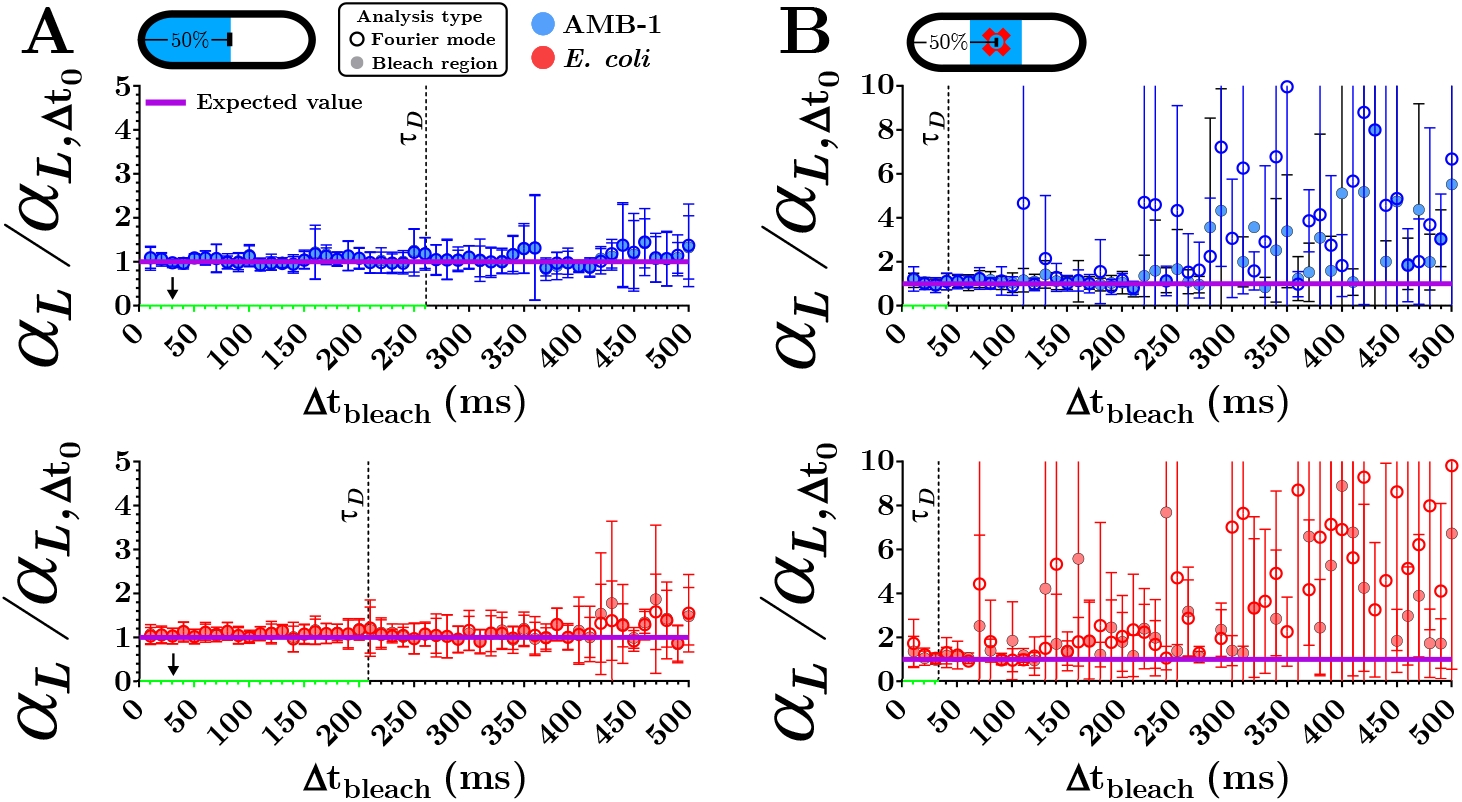
Dependence of recovery time on photobleaching step duration. Normalized recovery time *α*_*L*_ as a function of photobleaching step duration Δ*t*_bleach_ for (A) half-compartment bleach and (B) central region bleach (*R*_*ℓ*_ = 1 µm) simulations. *α*_*L*_ was normalized by its estimated value when 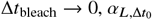. Simulations were mostly performed in the same conditions as those in Fig. 6: *L*_*T*_ = 5 µm, *D* = 15 µm^2^ s^−1^, *L*_*C*_ *L* = 1 (*E. coli*, red symbols) or 1.12 (*P. magneticum*, blue symbols), and frame acquisition interval Δ*t*_frame_ = 30 ms. In this case, however, the value of the fluorophore lifetime was kept constant, *λ*_bleach_ = 2 ms, although Δ*t*_bleach_ was varied. In each panel, the value of 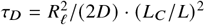 is indicated by a vertical dashed line. Each data point is the mean ± SD of 10 simulation replicates. Green-shaded regions demarcate the conditions in which the halo effect can be neglected. In (A), the conditions used in the accompanying microscopy experiments are marked with arrows.

For half-compartment bleach simulations, the normalized recovery time *α*_*L*_ varies very little over the explored range of photobleaching durations (Fig. 7A). The relatively large size of the bleach region in this case (*R*_*ℓ*_ = 2.5 µm) means that τ_*D*_ is also large (261 ms for AMB-1 and 208 ms for *E. coli*), such that at most Δ*t*_bleach_ = 2.5τ_*D*_ in the explored range. Even when Δ*t*_bleach_ > τ_*D*_, the recovery time remains constant, although the precision with which it is measured decreases. This behavior can be related to the weak dependence of the recovery time on bleach region size in half-compartment FRAP experiments (Fig. 6A). In contrast, using a small (*R*_*ℓ*_ = 1 µm) centrally located bleach region leads to much shorter values for τ_*D*_ (42 ms for AMB-1 and 33 ms for *E. coli*), and when Δ*t*_bleach_ becomes larger than a few τ*D*, both the average value of *α*_*L*_ the precision with which it is estimated start increasing (Fig. 7B). This observation is consistent with the sensitivity of the recovery time on central bleach region size (Fig. 6B). The influence of the frame acquisition interval post-bleach (fixed at Δ*t*_frame_ = 30 ms for the simulations shown in Fig. 7 to match the value used in the accompanying FRAP experiments) was also investigated (see Supplementary Material), confirming that when Δ*t*_bleach_ > τ_*D*_ the precision on recovery time measurements starts decreasing and this issue is exacerbated by slower frame acquisition.

Overall, our simulations confirm that half-compartment FRAP assays are more robust against variations in photobleaching duration than typical central region FRAP assays. This conclusion holds for cylindrical and helical geometries, and for both Fourier mode analysis and bleach region analysis.

### 3.3 Determination of the diffusion coefficient of mNG in AMB-1

#### 3.3.1 Implementation of half-compartment FRAP in AMB-1

As a first application of the proposed FRAP framework in helical bacteria, we compared the diffusion of the small soluble and constitutively monomeric fluorescent protein mNG in *E. coli* and AMB-1. mNG has a brightness roughly triple that of the Enhanced Green Fluorescent Protein (EGFP) and a very short maturation time (47–49). These desirable properties allowed working at low fluorescent protein concentrations (using a low copy plasmid under the *lac* promoter) which minimized the metabolic stress of non-native protein expression in the sensitive AMB-1 cells, and during early log phase when the cells are the healthiest and better able to withstand any perturbations during sample preparation and imaging. We assayed the effect of varying this chronic stress (using a low copy plasmid under an inducible *tac* promoter by inducing AMB-1 at lag phase post subculturing) caused by protein expression on the diffusion coefficient of mNG. We used the more robust half-compartment FRAP assays for all experiments.

For FRAP experiments, cells were immobilized on coverslips coated with Poly-L-Lysine. Cellbrite® Fix 640 membrane stain was used to identify cells with a compromised plasma membrane that resulted in stain internalization (Fig. 8). A total of 258 AMB-1 and 81 *E. coli* cells with an intact plasma membrane were selected for FRAP experiments. The result of representative half-compartment FRAP experiments, for both *E. coli* and AMB-1 cells of different lengths, as well as their analysis according to Fourier mode and bleach region analysis, are shown in Fig. 9A-D. Approximation of the fluorescence recovery by a single exponential decay was satisfactory for all cells studied and for both bleach region analysis and Fourier mode analysis. However, in contrast with simulations, a small difference in the fluorescence of bleached and non-bleached compartments was sometimes observed, before photobleaching and after recovery (e.g. Fig. 9A), which we attribute to a slight inclination of some cells with respect to the focal plane. This resulted in a non-zero value of Δ*I*_*∞*_, but did not preclude the extraction of the characteristic recovery time τ.

**Figure 8.**
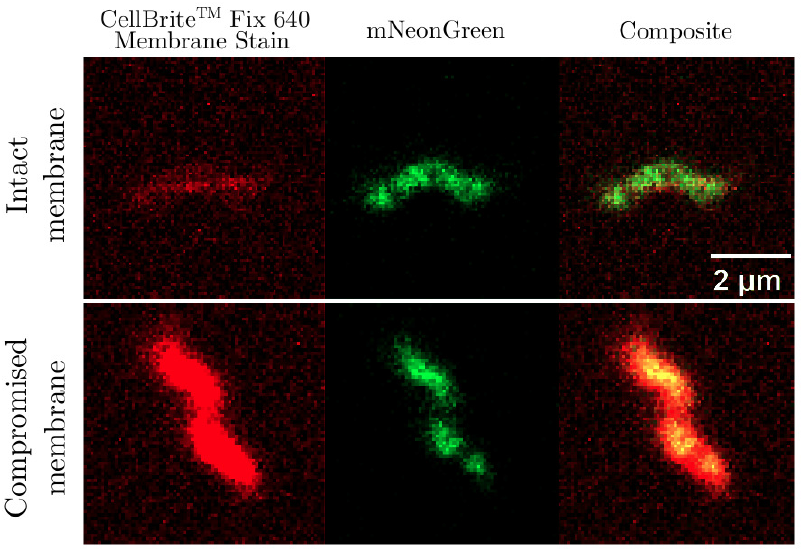
Identification of AMB-1 cells with a compromised cell membrane. Two-channel confocal images of AMB-1 cells expressing mNG and stained with Cellbrite® Fix 640 stain. The stain is only able to penetrate cells with a compromised cell membrane.

**Figure 9.**
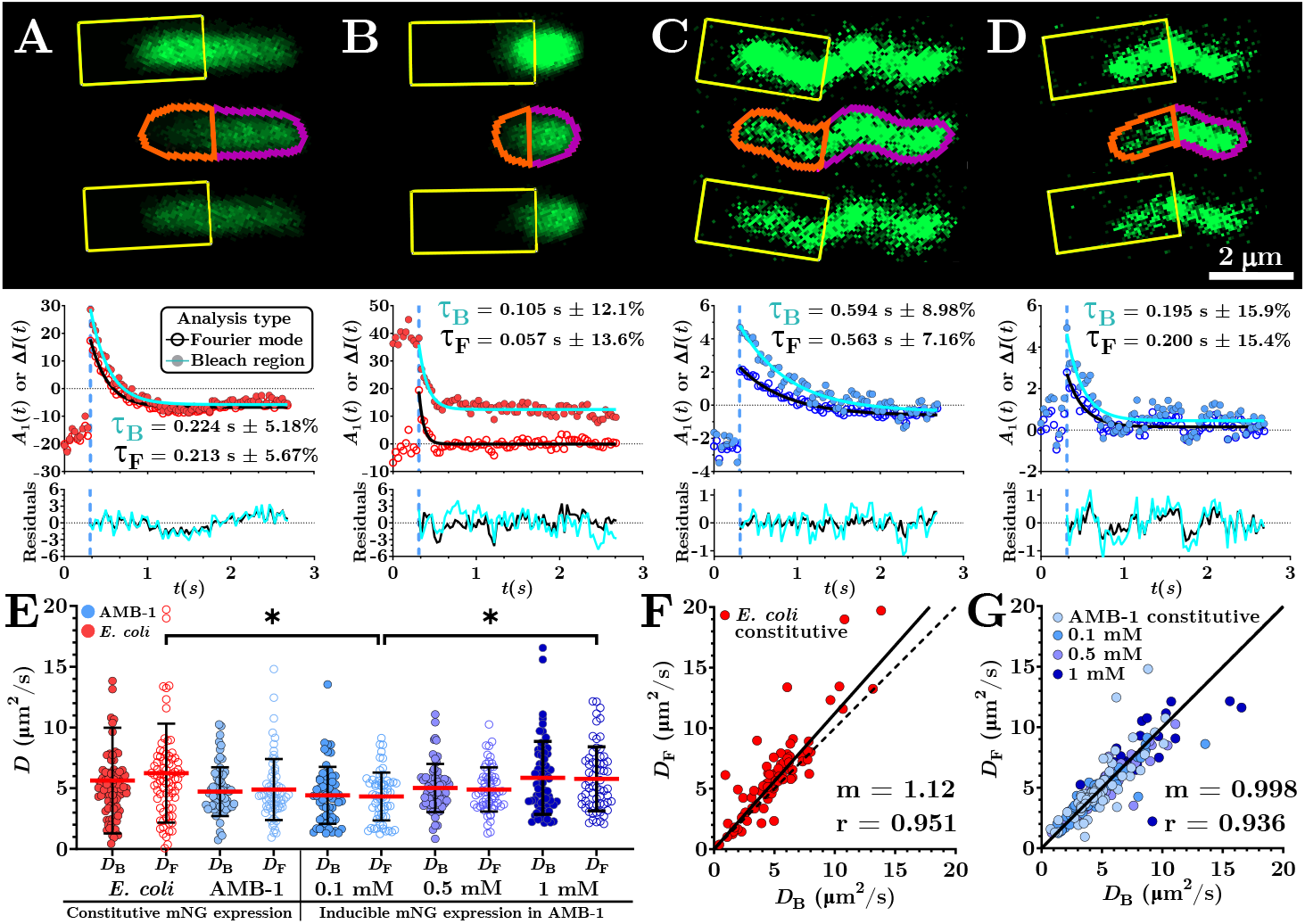
Diffusion of mNG within the cytoplasm of *E. coli* and AMB-1. Representative examples of half-compartment FRAP experiments conducted on (A,B) *E. coli* cells and (C,D) AMB-1 cells of different lengths expressing mNG. Upper panel: Confocal image of cells before, just after, and long after photobleaching. Lower panel: Recovery curves obtained using either bleach region or Fourier mode analysis. (E) Distribution of diffusion coefficients obtained for mNG in *E. coli* cells (n = 81) and AMB-1 cells (purely constitutive mNG expression, n = 67; inducible pseudo-constitutive expression with 0.1 mM IPTG. n = 63; 0.5 mM IPTG, n = 64; 1 mM IPTG, n = 64). (F,G) Correlation between the diffusion coefficients derived for each cell from bleach region analysis (*D*_*B*_) and Fourier mode analysis (*D*_*F*_), for either *E. coli* (F) or AMB-1 (G). Continuous lines are linear fits of the data, and *m* is the slope.

#### 3.3.2 Bleach region and Fourier mode analyses return comparable values for the diffusion coefficient of mNG

For each cell, Eq. 5 was used to calculate the cytoplasmic diffusion coefficient of mNG twice, first from bleach region analysis (*D*_*B*_) and then from Fourier mode analysis (*D*_*F*_), taking into account the cell’s unique size and geometry. The distribution of values obtained for *D*_*B*_ and *D*_*F*_ are shown side by side in Fig. 9E for both species and in the case of AMB-1 for different induction and expression regimes. Average values are listed in Table 1. For *E. coli*, 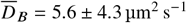 and 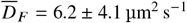, with no statistically significant difference between the values obtained by these two different methods (*p* = 0.9956). For AMB-1, the only statistically significant difference was was observed when comparing the means of *D*_*F*_ for the 0.1 mM induction condition and the 1 mM induction condition (*p* = 0.0211). All other comparisons between analysis method and expression regime showed no significant differences (all *p* ≥ 0.0752 with 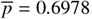, Brown-Forsythe one-way ANOVA with Games-Howell multiple comparisons test). Across species, the only significant difference was observed between 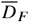 in *E. coli* and in AMB-1 at 0.1 mM IPTG (*p* = 0.0111). Regardless of species and conditions, we observed a very strong correlation between the values of *D* obtained using either analysis method (Fig. 9F,G). Overall, this confirms that using Eq. 5 to calculate *D* appropriately takes into account the small systematic difference in the value of the recovery time obtained from both analysis methods.

**Table 1:** mNG diffusion coefficients measured in AMB-1 and *E. coli* for various expression regimes using either Fourier mode analysis (*D*_*F*_) or bleach region analysis (*D*_*B*_). The weighted average diffusion coefficient (*D*_*IVW*_) and average fluorescence intensity pre-bleach (*I*) are also shown. Mean ± SD.

| | Constitutive<br>mNG Expression | | Inducible<br>mNG expression in AMB-1<br>(Pseudo-constitutive) | | | Pooled<br>AMB-1<br>($n = 258$) |
| --- | --- | --- | --- | --- | --- | --- |
| | *E. coli*<br>($n = 81$) | AMB-1<br>($n = 67$) | 0.1 mM<br>($n = 63$) | 0.5 mM<br>($n = 64$) | 1 mM<br>($n = 64$) | |
| $\overline{D}_F \pm \sigma$ ($\mu\text{m}^2 \text{s}^{-1}$) | $6.2 \pm 4.1$ | $4.9 \pm 2.5$ | $4.3 \pm 2.0$ | $4.9 \pm 1.8$ | $5.8 \pm 2.6$ | $5.0 \pm 2.3$ |
| $\overline{D}_B \pm \sigma$ ($\mu\text{m}^2 \text{s}^{-1}$) | $5.6 \pm 4.3$ | $4.7 \pm 2.0$ | $4.4 \pm 2.3$ | $5.0 \pm 2.0$ | $5.9 \pm 3.0$ | $5.0 \pm 2.4$ |
| $\overline{D}_{IVW} \pm \sigma$ ($\mu\text{m}^2 \text{s}^{-1}$) | $5.4 \pm 3.4$ | $4.7 \pm 2.0$ | $4.3 \pm 2.1$ | $4.9 \pm 1.8$ | $5.7 \pm 2.7$ | $4.9 \pm 2.2$ |
| $\bar{I} \pm \sigma$ (a.u.) | $61 \pm 34$ | $12 \pm 3$ | $52 \pm 14$ | $25 \pm 15$ | $22 \pm 7$ | $28 \pm 18$ |

#### 3.3.3 The diffusion coefficient of mNG is similar in *E. coli* and AMB-1 and largely independent of expression level

A meta-analytical value for *D*, combining the values obtained for both types of analysis, was obtained using Inverse-Variance Weighting (IVW) that is reported here as *D*_*IVW*_. The weighted average of *D*_*B*_ and *D*_*F*_ was calculated for each cell using 1/ *σ*^2^ where *σ*_*B*_ and *σ*_*F*_ are the one SD error for *D* calculated from the fitting error on τ for each analysis type (see Supplementary Material for details on fitting errors). There was no statistically significant difference in the mean value of *D*_*IVW*_ between both species (*p* ≥ 0.0970 with 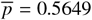), while the 0.1 and 1 mM IPTG condition had means that were significantly different (*p* = 0.0117) (Fig. 10A,B). While the means are not significantly different, the SDs for *D*_*IVW*_ were found to be significantly smaller for all AMB-1 conditions compared to *E. coli* (unpaired Welch’s t-tests *p* < 0.0498; F-test values between 1.613 and 3.417). In contrast, there was no statistically significant difference in the SD for the different conditions that the AMB-1 were subjected to. When pooling together all 258 AMB-1 cells to ascertain a *D* value representative of this helical species across a large spectrum of protein expression capabilities, we find *D*_*IVW*_ = 4.9 ± 2.2 µm^2^ s^−1^. Regardless of the expression levels in the helical AMB-1 cells, there is no significant difference in the mean compared to *E. coli* (*p* = 0.1718) but the variance is significantly different (*p* < 0.0001) (Fig. 10B). Thus, interestingly, assuming that the diffusion of mNG is reflective of cytoplasmic viscosity, AMB-1 cells have a variance in cytoplasmic viscosity that is 2.4× smaller than *E. coli*.

**Figure 10.**
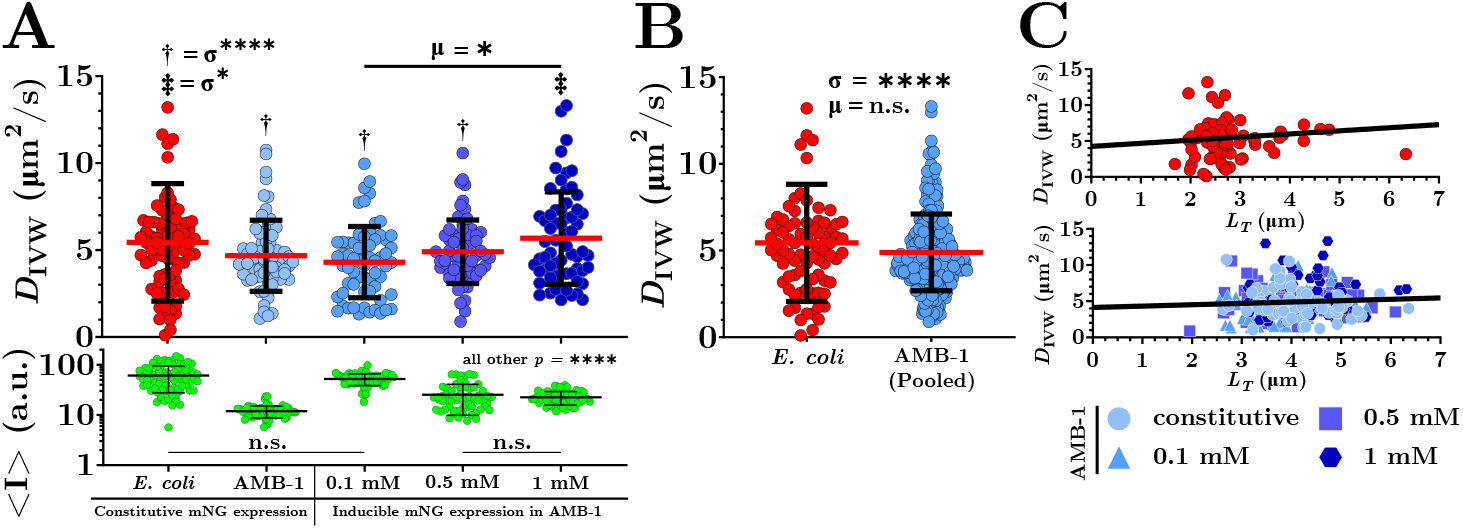
Dependence of mNG diffusion on expression level, cell species and cell length. (A) Weighted-averaged mNG diffusion coefficient, *D*_*IVW*_, obtained for constitutive expression or induced pseudo-constitutive expression with different IPTG concentrations. Single daggers denote AMB-1 conditions with very significantly different (*p* < 0.0001) variances compared to *E. coli* and double daggers denote a lower degree of significance (*p* = 0.0498). Lower panel: Average mNG fluorescence intensity recorded for each cell pre-bleach. Aside from the two noted exceptions, all distributions shown in this panel have significantly different means (*p* < 0.0001). (B) Distribution of mNG diffusion coefficients observed in *E. coli* and AMB-1 cells (regardless of expression level). (C) Diffusion coefficient as a function of cell length for *E. coli* (upper panel) and AMB-1 (lower panel).

Assuming that the viscosity of the prokaryotic cytoplasm is the same for cells of different lengths, we expect the value of *D* to be independent of *L*_*T*_. A linear regression analysis was performed on *D*_*IVW*_ for both cell types (Fig. 10C). The slopes of this regression were not significantly different from 0 for neither *E. coli* (*m* = 0.43, *p* = 0.4086) nor AMB-1 (*m* = 0.19, *p* = 0.3050). This confirms that in the 2 − 6 µm cell length range explored in these experiments Eq. 5 correctly takes into account the variation of recovery time with cell length, allowing recovery of the correct value of *D* regardless of the cellular geometry, and that the cytoplasmic viscosity was mostly unaffected by the length of either bacteria.

## 4 DISCUSSION

### 4.1 Cytoplasmic microviscosity of AMB-1

Much of what is known about cytoplasmic viscosity derives from measurements of the diffusion of small monomeric *β*-barrel fluorescent proteins such as GFPs. Their diffusion coefficient, 80 − 100 µm^2^ s^−1^ in aqueous solutions (50–53), is reduced to 20 − 30 µm^2^ s^−1^ when expressed in the cytoplasm of eukaryotic cells (51, 54), and further to 3 − 20 µm^2^ s^−1^ when expressed in bacteria (5, 7, 8, 21, 22, 55–58). For *E. coli* specifically, when cells are grown in rich LB medium, the average diffusion coefficient of EGFP is in the range 3 − 8 µm^2^ s^−1^ (5, 7, 22, 55–57). This value is very sensitive to growth conditions - increasing the osmolality of the growth medium or submitting cells to osmotic shock reduces the cytoplasmic water fraction and decreases the diffusion coefficient of EGFP (7, 8, 55, 57). A similar picture emerges for other bacteria. In the rod-shaped gram-negative pathogenic strain *Pseudomonas aeruginosa*, mCherry has an average diffusion coefficient of 3.7 µm^2^ s^−1^ (59). In spherical *Lactococcus lactis*, a Gram-positive diplococcal strain used in dairy fermentation, the diffusion coefficient of EGFP is ~ 7 µm^2^ s^−1^ (58). Thus the viscosity experienced by small proteins (microviscosity) in the cytoplasm of bacterial species studied to date is roughly 5 − 30 larger than in water, and 1.5 − 10 times larger than in the eukaryotic cytoplasm. Yet, microviscosity measurements in prokaryotes have largely been limited to spherical and rod-shaped cells, whereas prokaryotic cells display a wide range of morphologies (e.g. spherical or ellipsoidal cocci, rod-shaped bacilli, filamentous actinomycetes, helical vibrios (60)).

Here, we specifically developed an experimental and data analysis FRAP workflow to investigate the cytoplasmic viscosity of cells with helical, spirillum-shaped morphologies, the third most common among prokaryotes. We used the freshwater magnetotactic bacterium AMB-1, originally isolated from a pond in Japan (61), as a representative helical prokaryote, and studied the diffusion of the small fluorescent protein, mNG in this species. mNG has superior fluorescence properties while retaining the same characteristic *β*-barrel structure and hydrodynamic radius as other fluorescent proteins. Its diffusion coefficient in aqueous solution has been reported to be 97 ± 9 µm^2^ s^−1^ in a phosphate buffer at *T* = 28 °C (62) (and thus 87 ± 8 µm^2^ s^−1^ at *T* = 23 °C). To meaningfully compare the cytoplasmic viscosity of AMB-1 to that of E. coli, we grew each bacterial species in osmotic conditions reflective of their natural environment, 85 mOsm L^−1^ for AMB-1 and 300 mOsm L^−1^ for *E. coli*. In these conditions, the average diffusion coefficient of mNG is remarkably similar in these two organisms: 4.9 µm^2^ s^−1^ (*σ* = 2.2) in AMB-1 and 5.4 µm^2^ s^−1^ (*σ* = 3.4) in *E. coli*. Our measurements indicate that in normal growth conditions and at *T* = 23 °C, the microviscosity of the AMB-1 cytoplasm is on average 16.5 ± 9 mPa s, about 20 times larger than that of water at that same temperature. Remarkably, when both species are grown in their natural osmotic conditions, the cytoplasmic microviscosity of AMB-1 is, within experimental error, indistinguishable from that of *E. coli*.

MTBs (magnetotactic bacteria) such as AMB-1 are unique among prokaryotes in their ability to produce magnetosomes - membrane-enclosed organelles that each house a magnetic nanocrystal (63–65). While AMB-1 is a Gram-negative flagellated bacteria with length and diameter comparable to those of *E. coli*, these two species differ markedly in terms of cellular geometry (helical for AMB-1 versus cylindrical for *E. coli*), habitat (aquatic and low osmolality for AMB-1 versus gut-dwelling and high osmolality for *E. coli*), and degree of specialization (highly specialized with strict oxygen requirements for AMB-1 versus metabolically versatile for *E. coli* which can thrive in a wide range of environments). Notably, doubling times are very different in these two species: 7 − 8 h for AMB-1 in the regular growth conditions used in this study (see Supplementary Materials) and ~4 h under optimal conditions (66), versus as short as 20 min for *E. coli* - representative of the wide range of doubling times observed in prokaryotes. Thus, our results show that the average growth rate of a prokaryote is not directly related to its cytoplasmic microviscosity. Doubling times must be controlled by other factors unique to their respective proteomes for reasons specific to the organism. In AMB-1, for example, the cell cycle might be linked to magnetosome biomineralization, which takes several hours (67–70).

Exposing a specific prokaryote to conditions that induce overcrowding induces both an acute increase in cytoplasmic microviscosity and a decrease in growth rate. In *E. coli*, increasing growth media osmolarity above native levels correlates with a decrease in GFP diffusion coefficient and an increase in cell doubling time (8). A slowing of growth rates in increasingly hypertonic solutions is also observed in *Magnetospirillum gryphiswaldense*, a MTB species closely related to AMB-1 (71). Although our study does not directly address the question of crowding, the slight differences in diffusion coefficients we observe for various mNG expression regimes in AMB-1 might be explained in the light of the correlation between crowding and microviscosity. First we note that increasing the IPTG induction level increased mNG concentration only up to a certain point after which it started decreasing (Fig. 10A), a phenomenon well documented in *E. coli* when using inducible systems. The lower *D* (higher microviscosity) observed for 0.1 mM IPTG could thus be due to a slight over-crowding caused by the high mNG concentration obtained in these conditions. In contrast, the harsher pseudo-constitutive induction regimes (0.5 mM and 1 mM IPTG) may have selected for cells that have lower basal rates of expression, capable of surviving under these more metabolically demanding forced expression conditions early on in their growth cycle. Such cells may be inherently less crowded, resulting in a shift toward what may be the optimal microviscosity for this species, suggesting biological regulation.

More broadly, our microviscosity measurements, the first in a helical species to our knowledge, support the hypothesis that cytoplasmic crowding and therefore microviscosity might be similar across prokaryotes when grown in favorable conditions. This optimal crowding could be the right balance between a high enough concentration of different solutes in the cell and a low enough microviscosity to ensure sufficient intracellular protein dynamics and metabolism (72). This is potential evidence for convergent evolution in prokaryotes across a range of microbial habitats with very different osmotic conditions. Other more commonly studied strains of helical bacteria, such as *Helicobacter pylori, Campylobacter jejuni*, or *Borrelia burgdorferi*, should be investigated to determine if other spirillum-shaped bacteria have similar intracellular dynamics.

### 4.2 Variability in cytoplasmic microviscosity

We find that *E. coli* has a high variance in cytoplasmic microviscosity, with 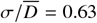, a value comparable to previously reported ones (7). This relative dispersion is larger than the 5 − 15% relative error on individual diffusion coefficient measurements, and therefore reflects real biological variations within the cell population. The relative dispersion in cytoplasmic microviscosity for AMB-1, 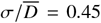, is slightly but significantly smaller than in *E. coli*, indicating lower cell-to-cell variability in microviscosity.

A high variability in cytoplasmic microviscosity can be expected to correlate with the capacity of a cell population to withstand and adapt to various sources of natural selection, for example, sudden changes in osmolarity, creating harsh hypotonic or hypertonic environments. The greater the chance of finding individuals that exhibit crowding and intracellular dynamics at either extreme, the more resilient that population will be to osmotic shock. *E. coli* is known to be a very robust organism, capable of withstanding large variations in salt concentrations. The strain *E. coli* K-12, for example, can sustain osmotic changes as high as + 1300 mOsm L^−1^ (73). In contrast, AMB-1, like other magnetospirilla, require a very specific environment to thrive. In fact, many magnetospirilla cannot be kept in cultures for prolonged periods. The salt tolerance for the closely related strain *Magnetospirillum gryphiswaldense* is ≤ + 80 mOsm L^−1^, with population growth slowing down above this value (71). Furthermore, interestingly, we can see that the AMB-1 populations in the various expression regimes have very tight distributions for the expression (using intensity as an analog) and have very small variance, whereas the *E. coli* have very high variance for the expression. This reiterates the fact that AMB-1 have very strict metabolisms, and any large disturbances, namely higher, forceful, and chronic induction conditions, will result in selecting for populations that express less, whereas *E. coli* have very expansive expression profiles, allowing them to survive in various environments and conditions.

### 4.3 Practical implications for the implementation of FRAP in bacteria and small compartments

The quantitative interpretation of FRAP experiments is challenging when bleaching is performed with arbitrary patterns, within small compartments, or in inhomogeneous media. Even in the case of a freely diffusing fluorophore these conditions generally preclude the derivation of simple analytical expression for *D*. In such cases, numerical simulations provide a powerful framework to model and guide the analysis of FRAP experiments (4, 74–76). Here, we developed FRAP simulations specifically tailored to diffusion within helical cells, allowing a systematic investigation of the factors that might influence the characteristic recovery time extracted from FRAP experiments, τ. This approach led to the simulation-derived phenomenological relationship linking *D* and τ captured in Eq. 5. The application of this analytical framework was applied systematically to the raw microscopy data with PyImageJ providing a useful bridge between Fiji’s image-processing capabilities and Python-based data analysis workflows.

A practical outcome of this work is the identification of experimental conditions under which the relationship between *D* and τ becomes largely independent of experimental FRAP parameters. Our simulations highlight that bleaching half of the cell provides a straightforward and robust FRAP strategy for studying diffusion in bacteria, and more generally, in small compartments. It results in fluorescence recoveries appropriately described by single-exponential dynamics (Fig. 3B,D), with a characteristic decay time that is insensitive to both the exact length of the bleached region (Fig. 6A) and the precise photobleaching step duration (Fig. 7A). This robustness allows for flexibility when choosing microscopy settings, and the possibility to use relatively large bleach regions, as well as longer bleach times and slower acquisition rates than when using smaller centrally-located bleach regions. Importantly, the use of a relatively large bleach region means that studying fast-diffusing proteins is possible. The advantages of half-compartment bleaching are generally relevant to FRAP studies in small compartments, including spherical compartments. Nonetheless, targeted bleaching of an inner section of the cell may still be advantageous in specific situations, for example to probe protein binding to a localized magnetosome cluster in AMB-1.

In half-compartment FRAP experiments performed in helical cells, the relevant length scale is the cell contour length, *L*_*C*_. To first order, in an elongated compartment fluorescence recovery is dominated by the decay of the first Fourier mode along the cell backbone, yielding the relationship 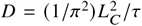. However, corrections are required for compartments with aspect ratios below ~ 10, a regime that encompasses most common types of bacterial cells. The magnitude of this correction can vary substantially even among cells of the same species, hence the importance of taking into account aspect ratio on a cell-by-cell basis. In addition, the method used to analyze the FRAP data introduces systematic differences in the measured values for τ (Fig. 5), because different types of analysis capture different combinations of relaxation modes. Fourier mode analysis, by isolating a single relaxation mode along the cell long axis, allows, in principle, the robust determination of a well-defined relaxation time. However, it is not well-adapted to the analysis of cells with small aspect ratios and a less well-defined one-dimensional fluorescence profile. Bleached region analysis, on the other hand, performs consistently for cells of all aspect ratios. However, as integrating the fluorescence over the bleached region combines features from a number of modes, the apparent relaxation time obtained from the single-exponential fit of the bleach region fluorescence decay is very sensitive to the time range and time resolution of the recorded post-bleach data. The general relationship between *D* and τ obtained from our simulations (Eq. 5) accounts for differences in cell geometry and analysis method via a simple change in parameters and enables the determination of diffusion coefficients for helical cells of arbitrary length, thickness, and helical parameters, using either bleach-region or Fourier-mode analysis. We recommend using bleach region analysis for cells with *L*_*T*_ /*d* ≤ 2, and Fourier mode analysis for those with *L*_*T*_ / *d* ≥ 10. For typical bacterial cells with *L*_*T*_ / *d* in the range 2 − 10, where these two methods give comparable performance, using both of them and calculating a weighted-average as done in this study might represent the best solution.

Beyond freely diffusing proteins in the cytoplasm of helical bacteria, this framework is readily applicable to a range of other systems. In particular, the closed-form expression derived for *D* can be applied to spherical compartments, which is highly relevant for studies of biomolecular condensates, as well as for larger cellular structures such as yeast cells or nuclei. Our simulation-based approach could also be easily adapted to describe diffusion in a bacterial membrane or periplasmic space, diffusion across porous or membraneless boundaries, different types of anomalous diffusion, and diffusion of molecules interacting with cellular structures. More generally, our FRAP methodology can be extended to cells with other elongated or complex morphologies, including shapes that can be generated from a one-dimensional backbone and, with additional effort, to non-rigid geometries. For example, diffusion measurements could be performed in Y-shaped Bifidobacteria, box-shaped Haloquadrata, or club-shaped Corynebacteria observed during specific growth phases.

## Supporting information

Osmolarity calculation worksheet

Supplemental video 1: Simulation animation

## Acknowledgments

FRAP experiments were conducted at the McMaster Centre for Advanced Light Microscopy (CALM).

TEM experiments were conducted at the Canadian Centre for Electron Microscopy (CCEM), also at McMaster University.

## Author contributions

S.S. and C.F. conceived the study and designed the research. S.S. designed and performed all the molecular cloning, bacterial transformations, microscopy, programming, simulations and data analysis. C.F. performed analytical calculations. S.S. and C.F. wrote and edited the manuscript. S.S. prepared visualizations. C.F. supervised the project and acquired funding.

## Data availability

Raw data are available through McMaster Dataverse, a collection within Borealis, the Canadian Dataverse Repository as follows: Shariful, Sakib; Fradin, Cécile, 2026, “Replication Data for: Experimental and simulated FRAP for the quantitative determination of protein diffusion in helical cells”, https://doi.org/10.5683/SP3/D7SBYN, Borealis, V1. The simulation and automated FRAP data analysis scripts are available on GitHub (here) and archived on Zenodo (here). The DNA sequences of the plasmids can be found on GenBank for the low expression plasmid pRK415-mNG (constitutive) with Accession #: PX514998 and for the pRK415-mNG (inducible) plasmid with Accession #: PX911503. The cloning strains of *E. coli* harbouring these plasmids are available through Addgene (plasmid #248204 RRID: Addgene 248204 and plasmid #252211 RRID: Addgene 252211).

## Supplementary Material

This article has a supplementary information file.

