## Supplementary information for "Experimental and simulated FRAP for the quantitative determination of protein diffusion in helical cells"

#### 1 Optimal growth medium for AMB-1 cells

AMB-1 cells were cultured in four different Magnetic Spirillum Growth Media (MSGM) for comparison. MSGM-LN is from Le Nagard *et al. 2018* (originally referred to as "MSR-1 Media") [1]. MSGM is the ATCC Medium 1653: Revised Magnetic spirillum growth Medium. Modified MSGM (mMSGM) is from Furubayashi *et al. 2021* [2]. Enriched MSGM (EMSGM) is from Yang *et al. 2001* [3]. AMB-1 were seed cultured into these different media from frozen glycerol stock then each of these cultures was subcultured in that same medium in parallel. Time point zero was defined as the time immediately after subculturing. Data was collected every 8 hours within a 64 hour period with a 633 nm laser. Each medium was inoculated and then incubated under stationary conditions.

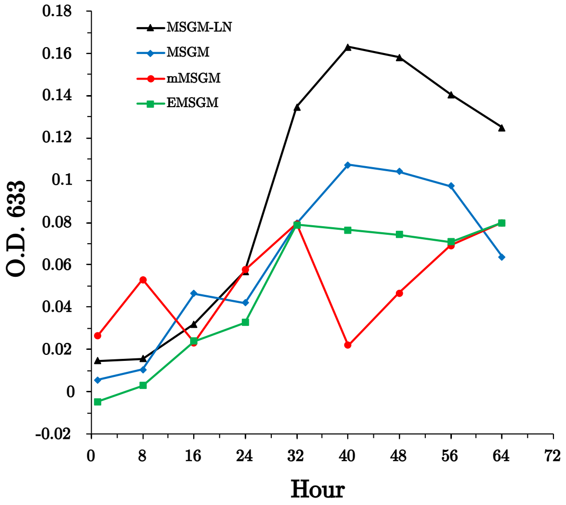


Figure S1: Growth curves for AMB-1 in different Magnetic Spirillum Growth Media reported in the literature. See text for details.

### 2 Construction of LacI repressor system via 5-fragment plasmid assembly

The following protocol is currently protected by US provisional (US 63/881,962) in the form of a molecular biology kit and cannot be used commercially without an agreement. It can be used freely for research purposes. The construction of over a dozen plasmids, ranging from small 2-fragment pUC19 ($\sim$ 3 kbp) assemblies all the way to large 5-7-fragment pRK415 assemblies ($\sim$ 14 kbp), has been achieved by this method. This protocol optimizes DNA inputs and outputs and was found to be very reproducible. The exact concentrations and compositions of buffers and enzymes are not stated as they are proprietary but can be substituted with any common lab formulations if being replicated by labs for research purposes.

#### 2.1 DNA Fragment generation and pre-assembly prep

1. DNA fragments with homologous terminal sequences can be generated by PCR (20 µL reactions), restriction enzyme digests, or chemical synthesis. They need to have at least 20 bp of overlap.

2. DNA fragments can be assessed on an agarose gel for target band specificity. Non-specific samples can be extracted from an agarose gel with a guanidine thiocyanate-based buffer with chaotropic salts. The gel weight (mg) to buffer volume (mL) ratio can range from 1:3 to 1:5.

3. This DNA solution is then loaded onto a column with a silica-based membrane via slow centrifugation at 3000 $\times$ g for 5 min for improved recovery.

   Note: All subsequent washes and elutions are conducted via centrifugation at 14,000 $\times$ g. All subsequent flow-through can be discarded, preferably by pipette aspiration.

4. The DNA bound to the column is washed twice with 200 µL of an ethanol-based buffer with guanidine-HCl to maintain chaotropic conditions to remove residual contaminants while preserving DNA binding to the silica membrane. (Wash buffers first sit on the membrane for 1 min to ensure maximal coverage of the silica).

5. A "dry spin" can be incorporated after removing the ethanol flowthrough. The DNA is then eluted into a 1.5 mL tube with 20-30 µL of nuclease-free water that has been warmed to 60-80°C via centrifugation for 1 min. Do not discard the spin-column with the silica-membrane.

6. The column is flushed with 3 column volumes of nuclease-free water to remove residual chaotropic salts and is reused as a secondary purification matrix.

7. The DNA solution from step 5 is then further purified by first mixing the DNA solution with an isopropanol-based binding buffer with a 5:1 to 2:1 ratio of buffer volume to DNA solution volume and loading the mixture onto the flushed spin-column from step 6 via centrifugation at 14,000 $\times$ g for 1 min. The flowthrough is then discarded.

8. The DNA is washed with 200 µL of an ethanol-based wash buffer twice and the flowthrough is discarded. (Wash buffers first sit on the membrane for 1 min to ensure maximal coverage of the silica membrane).

9. The spin column is centrifuged again with no buffer ("dry spin") to ensure that no residual buffer remains. The column is transfered to a 1.5 mL tube.

10. A 20 µL volume of pure nuclease-free water heated to 60-80°C is then pipetted and the entire spin column apparatus is incubated on a heat block (or water bath). The DNA is finally eluted via centrifugation at 14,000 $\times$ g for 1 min.

#### 2.2 DNA assembly

Standard isothermal DNA assembly is conducted by using a 2X master mix (contains dNTPs, proofreading DNA polymerase, 5' exonuclease, T4 DNA ligase, PEG 8000, NAD+, DTT, MgCl2, Tris-HCl to pH 7-8). The input DNA can be quantified with a microvolume spectrophotometer or fluorometer. The DNA fragments can be input in equimolar ratios (anything between 20 - 200 fmol per fragment) and mixed with the 2X master mix. The mixture is incubated at 50°C for 1 hour (optimally in a PCR machine/thermocycler).

#### 2.3 Post-assembly prep

The assembled plasmid DNA is then treated with 10 Units of DpnI in conjunction with an exonuclease or combination of exonucleases that exhibit ssDNA, dsDNA, 5' and 3' catalytic activity at 37°C for 1 hour. This combination of enzymes is capable of digesting any unreacted linear fragments or any template DNA down to the individual nucleotides that could hinder downstream applications such as bacterial transformation. The covalently closed plasmid DNA is spared from this enzymatic cocktail, thus, this mixture can be purified further with another round of spin-column purifications as described in step 3 to 10 for the purposes of isolating the plasmid construct of interest and to reduce the incidence of polymeric contaminants that can prevent the pDNA from crossing the membrane pores in chemically competent *E. coli.* For the purposes of transformation, the plasmid can be eluted with 20 µL of nuclease-free water warmed to 60-80°C and stored at -20°C for up to one year. Standard bacterial transformation protocols can be followed.

### 3 Sample preparation for FRAP experiments in bacteria

A visual set of instructions to assist in the sample preparation to conduct FRAP experiments on bacteria such as the one described in the main text is shown in Fig. S2. The membrane dye can be used to identify cells that have a compromised membrane.

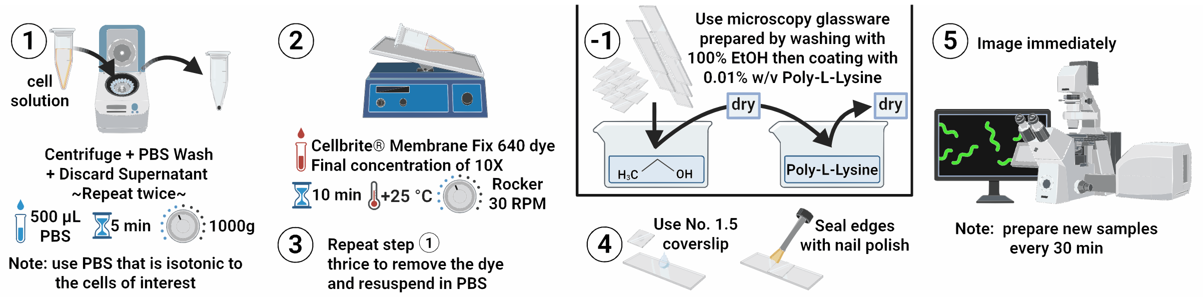


Figure S2: **Bacteria sample preparation for FRAP experiments**. Created in BioRender. Sakib, S. (2026) https://BioRender.com/2lcugsj.

### 4 Analytical estimate of bleach region fluorescence recovery for half-compartment FRAP experiments

This section includes a presentation of analytical derivations of the fluorescence recovery obtained in half-compartment FRAP experiments for two ideal limit cases (cylindrical and spherical compartments, infinitely short photobleaching step leading to bleach regions with sharp boundaries). The discussion of these ideal cases support the understanding of the simulation results presented in the main text.

#### 4.1 Cylindrical compartment

We first consider the case of a cylindrical compartment of length $L$ with flat ends. This case is relevant to FRAP experiments in very long cells with large aspect ratio, when spherical caps at the end of the cell become irrelevant.

##### 4.1.1 Decomposition of fluorescence profile in Fourier modes

Assuming a concentration gradient along the cylinder axis reduces the system to a one-dimensional fluorescence intensity profile, $i(x,t)$. For a single fluorophore species with diffusion coefficient $D$, $i(x,t)$ must obey the one-dimensional diffusion equation:

$$\frac{\partial i(x,t)}{\partial t} = D\frac{\partial^2 i(x,t)}{\partial x^2} \tag{S1}$$

and limit conditions: $\frac{\partial i(x,t)}{\partial x}|_{x=0} = 0$ and $\frac{\partial i(x,t)}{\partial x}|_{x=L} = 0$. The intensity profile can be decomposed into a Fourier series:

$$i(x,t) = \sum_{k=0}^{\infty} A_k(t)\cos(k\pi x/L). \tag{S2}$$

As each Fourier mode must separately obey the diffusion equation:

$$A_k(t) = A_{k,0}e^{-\frac{t}{L^2/(Dk^2\pi^2)}}. \tag{S3}$$

The amplitude of each mode can be determined from the initial profile:

$$A_{k,0} = \frac{2}{L}\int_0^L i(x,0)\cos(k\pi X/L)dx. \tag{S4}$$

##### 4.1.2 Time-dependent fluorescence profile

We consider FRAP experiments where a region of length $R_\ell$ is bleached, starting from the cylinder's leftmost extremity. The normalized intensity profile right after photobleaching is a step function, where $i(x,0) = 0$ for $x < R_\ell$ and 1 for $x > R_\ell$ (Fig. S3A).

The amplitude of each Fourier mode can then be explicitly calculated:

$$A_{0,0} = \frac{L - R_l}{L} \tag{S5}$$

and:

$$A_{k,0} = -2\frac{\sin(k\pi R_l/L)}{k\pi} \tag{S6}$$

for $k > 0$.

For a perfect half-bleach experiment ($R_\ell = L/2$), all even modes except for the $k = 0$ mode have an amplitude of zero (Fig. S3B). The dominant mode is the $k = 1$ mode, with relaxation time:

$$\tau_1 = \frac{L^2}{\pi^2 D}. \tag{S7}$$

Other non-zero modes, with relaxation time $\tau_k = \tau_1/k^2$, have diminishing amplitudes scaling with $1/k$.

As $R_\ell$ increases, the amplitudes of the $k = 0$ mode and that of odd modes decrease, as those of $k > 0$ even modes increase. While the $k = 1$ mode remains dominant, the next largest $k = 2$ mode progressively gain in importance (Fig. S3B).

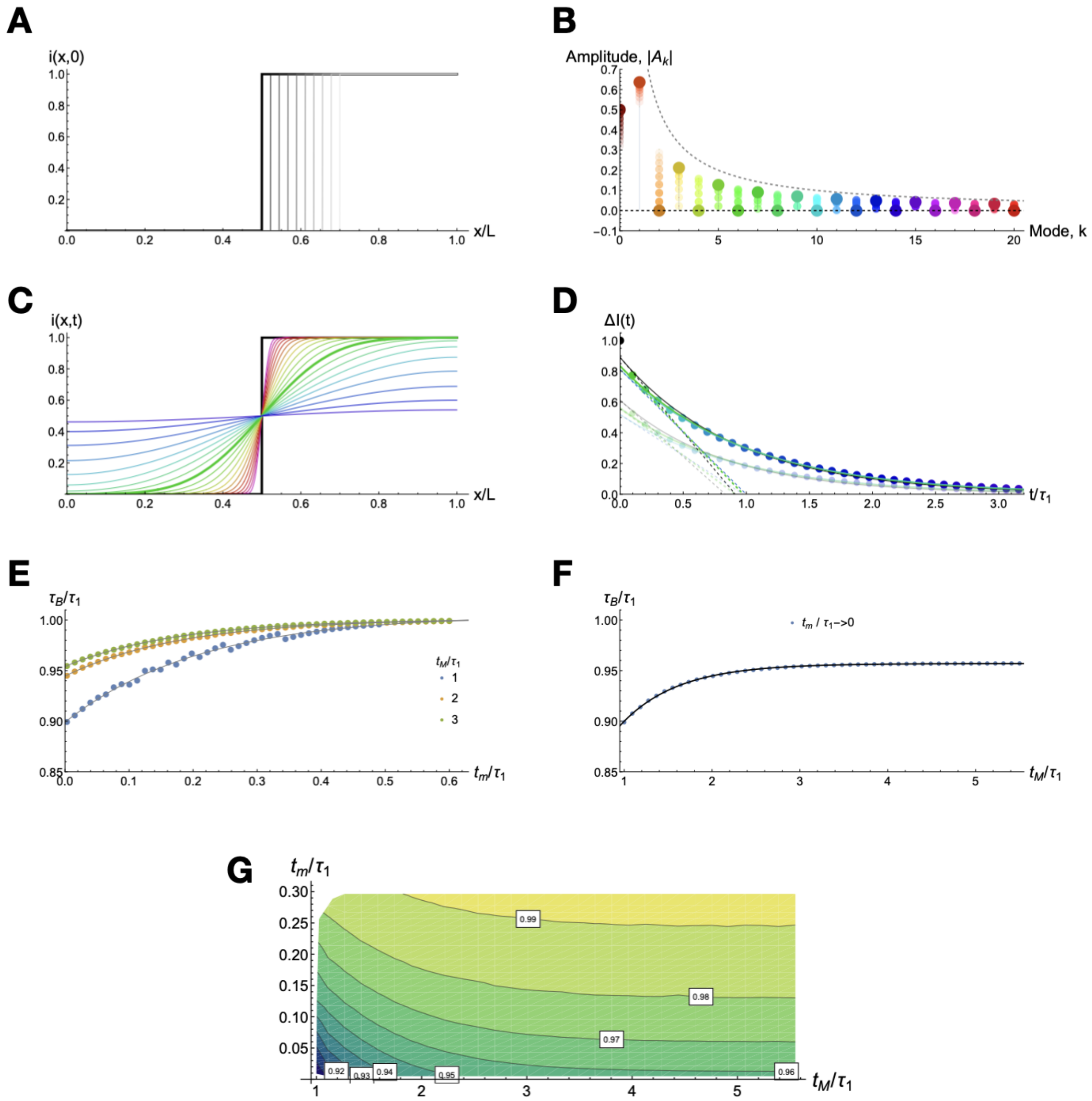


Figure S3: **Half-bleach FRAP in a cylindrical compartment.**. (A) Intensity profile right after photobleaching for instantaneous half-bleach (thick black line, $R_\ell = L/2$) or close to half bleach (thin grey lines, $R_\ell > L/2$) FRAP experiments. (B) Amplitude of the first 20 Fourier modes for the profiles shown in (A). Opaque dots correspond to the half-bleach profile, while smaller increasingly transparent dots correspond to profiles with increasing $R_\ell$. The amplitudes of the Fourier modes are in this case bounded by 0 and $1/k$ (black and gray dashed lines). (C) Intensity profiles after compartment half-bleach, represented immediately after photobleaching ($t = 0$, thick black line) and for a series of logarithmically spaced time points ($t_n = 0.0001 \times 1.52^n \times L^2/D$, coloured lines). The thick green line corresponds to $t_{11} = 0.01 \times L^2/D$. (D) Fluorescence decay calculated for half-bleach ($R_\ell = 0.5L$, large opaque circles) or close to half bleach ($R_\ell > 0.65L$, small transparent circles) FRAP experiments. Continuous lines show exponential fit of the data when using the full data range (black lines), when omitting the first point at $t = 0$ (green lines) or when omitting the first 5 points (blue lines). Dashed lines show the tangents to these fits at $t = 0$. The intersection of the dashed line with the x-axis corresponds to the value of $\pi^2 \tau_B D/L^2 = \tau_B/\tau_1 = \pi^2 \alpha_L$ returned by the fit. (E) Apparent relaxation time $\tau_B$ (normalized by $\tau_1$) obtained from the fit of fluorescence relaxation curves (such as as the ones shown in (D)) between $t_m$ and $t_M$. The data is plotted as a function of frame interval ($t_m$, which is also the first data point included for the fit), for three different series each corresponding to a different value of the total experiment time ($t_M$). Lines show exponential fit of the data. (F) Value of $\tau_B/\tau_1$ in the limit where $t_m/\tau_1 \to 0$, as extrapolated from exponential fits such as the ones shown in (E). The line shows an exponential fit of the data. (G) Contour plot showing $\tau_B/\tau_1$ as a function of both $t_m/\tau_1$ and $t_M/\tau_1$.

The time-dependent fluorescence profile is given by:

$$i(x,t) = \frac{L-R_l}{L} - 2\sum_{k=1}^{\infty} \frac{\sin(k\pi R_l/L)}{k\pi} \cos(k\pi x/L) e^{-\frac{k^2\pi^2 D}{L^2}t}. \tag{S8}$$

For a perfect half-bleach experiment, this profile very quickly relaxes to a shape close to a sinusoidal (due to the quick decay of modes with $k > 1$), then more slowly approaches a uniform profile (due to the slower decay of the $k = 1$ mode) (Fig. S3C). The thick green line in Fig. S3C corresponds to a time $0.01 \times L^2/D \simeq 0.1\tau_1$, which for cylinders of length $L = 2 - 5$ $\mu m$ and fluorophores with $D = 5\,\mu\mathrm{m}^2\,\mathrm{s}^{-1}$ (standard experimental conditions for FRAP in bacteria) is $8 - 50$ millis (comparable to sampling time at which images are captured after photobleaching in a tyical FRAP experiment). This profile is thus representative of the first profile that might be captured just after photobleaching in a typical half-bleach FRAP experiment for a soluble cytoplasmic protein in a bacterial cell.

###### 4.1.3 Fluorescence recovery in bleach region

The average fluorescence in the bleached region can be calculated by integration:

$$I_b(t) = \frac{1}{R_\ell}\int_0^{R_l} i(x,t)dx = \frac{L-R_l}{L} - \frac{2L}{\pi^2 R_l}\sum_{k=1}^{\infty}\frac{1}{k^2}\sin^2(k\pi R_l/L)e^{-\frac{k^2\pi^2 D}{L^2}t}. \tag{S9}$$

Similarly for the non-bleached region:

$$I_n(t) = \frac{1}{L-R_\ell}\int_{R_l}^{L} i(x,t)dx = \frac{L-R_l}{L} + \frac{2L}{\pi^2 R_l}\sum_{k=1}^{\infty}\frac{1}{k^2}\sin^2(k\pi R_l/L)e^{-\frac{k^2\pi^2 D}{L^2}t}. \tag{S10}$$

Thus the difference between the average fluorescence in these two compartments is simply:

$$\Delta I(t) = I_n(t) - I_b(t) = \frac{4L}{\pi^2 R_l}\sum_{k=1}^{\infty}\frac{1}{k^2}\sin^2(k\pi R_l/L)e^{-\frac{k^2\pi^2 D}{L^2}t}, \tag{S11}$$

or:

$$\Delta I(t) = \frac{L}{R_l}\sum_{k=1}^{\infty} A_{k,0}^2 e^{-\frac{k^2\pi^2 D}{L^2}t}. \tag{S12}$$

Thus the fluorescence decay is a sum of exponential decays, strongly dominated by the first ($k = 1$) Fourier mode. This mode has a characteristic decay $\tau_1 = L^2/(\pi^2 D)$ which is longer than that of all the other ($k > 1$) Fourier modes. As a result, when fitting the bleach region fluorescence decay to a single-exponential function, the obtained apparent decay time $\tau_B$ should be equal or slightly shorter than $\tau_1$. In other words the normalized decay time $\alpha_L^B = \tau_B D/L^2$ should be slightly shorter than $1/\pi^2$, i.e. $\pi^2\alpha_L^B \lesssim 1$. Importantly, we also expect that the value of $\alpha_L^B$ will depend on the range of the fit and the sampling of the data. This is illustrated in Fig. S3D, which shows a sampling of the fluorescence decay, and several fits performed using a different starting point post-bleach - each of these fits return a slightly different value for $\pi^2\alpha_L^B$.

The first recovery data point obtained in a FRAP experiment is captured at a time $t_m > 0$ post-bleach, where $t_m$ is the frame interval. As $t_m$ is increased and approaches $\tau_1$ (while keeping the total duration of the post-bleach observation, $t_M$, constant), the apparent recovery time approaches the value of $\tau_1$ (Fig. S3E). Similarly, as the experiment duration $t_M$ is increased well past $\tau_1$ (for a fixed value of $t_m$), the recovery time quickly increases to a plateau value (Fig. S3F, where this increase is shown for $t_m/\tau_1 \to 0$). In standard robust experimental conditions, when $t_m/\tau_1 < 100$, and $t_M/\tau_1 > 0$ we find that $\tau_B/\tau_1$ is comprised between 0.95 and 0.96 (Fig. S3F).

###### 4.1.4 Close to half-bleach FRAP experiments

If the bleach region is slightly larger or slightly smaller than exactly half the cell, even Fourier modes gain in importance, in particular mode $k = 2$. As a result, the recovered recovery time becomes shorter than in the case of a perfect half-bleach experiment (Fig. S3D). As a result the value of $\tau_B/\tau_1$ systematically decreases as $R_\ell$ deviates from the ideal $L/5$ value (Fig. S4A). In parallel, as $R_\ell$ deviates from $L/5$, the $k = 1$ mode becomes less dominant and the shape of the fluorescence decay becomes less well-approximated by a single exponential (Fig. S4B).

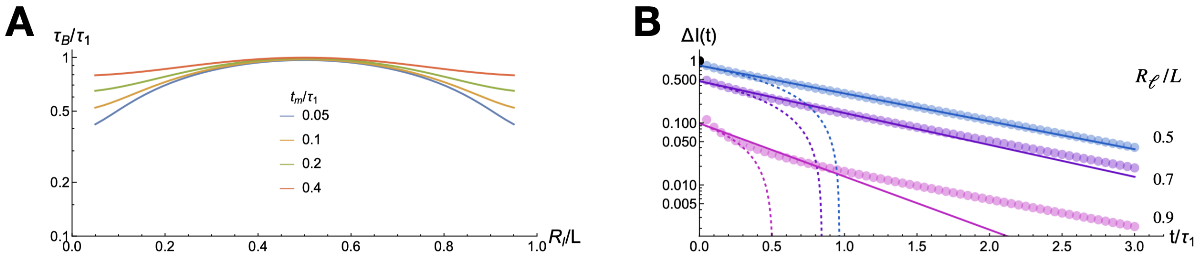


Figure S4: **Close to half-bleach FRAP in a cylindrical compartment.** (A) Normalized bleach region recovery time $\pi^2\alpha_L^B = \tau_B/\tau_1$ as a function of length of the bleach region $R_\ell$ relative to the total length of the compartment $L$. (B) Example of bleach region fluorescence decays plotted on a log-scale.

#### 4.2 Spherical compartment

We next consider the case of a spherical compartment of radius $R = L/2$. This case is relevant to FRAP experiments in spherical cocci, yeast cells, eukaryotic cell nuclei or spherical condensates.

##### 4.2.1 Decomposition of the fluorescence profile into spherical Bessel functions

For a spherical compartment, instead of a decomposition into Fourier modes, the fluorescence profile, $i(r,\theta,\phi)$, should be decomposed into spherical harmonics ($Y_{lm}(\theta,\phi)$) and spherical Bessel functions ($j_l(k_{ln}r)$):

$$i(r,\theta,\phi,t) = \sum_{l=0}^{\infty}\sum_{m=-l}^{+l}\sum_{n=1}^{\infty} B_{lmn}(t)Y_{lm}(\theta,\phi)j_l(k_{ln}r). \quad \text{(S13)}$$

In order for the no-flux boundary condition $\frac{\partial i(r,\theta,\phi)}{\partial r}|_{r=R} = 0$ to be respected, the radial wavenumber $k_{ln}$ must obey: $\frac{dj_l(k_{ln}r)}{dr}|_{r=R} = 0$, thus $k_{ln}R$ is the nth zero of $j_l'$.

As before each mode must fulfill the diffusion equation, leading to:

$$B_{lmn}(t) = B_{lmn,0}e^{-Dk_{ln}^2 t}. \quad \text{(S14)}$$

The amplitude of the modes with $n > 0$ can be calculated from the initial fluorescence profile $i_0(r,\theta,\phi)$ as:

$$B_{lmn,0} = \frac{\int_0^R i_{lm,0}(r)j_l(k_{ln}r)r^2dr}{\int_0^R j_l^2(k_{ln}r)r^2dr}, \quad \text{(S15)}$$

and:

$$i_{lm,0}(r) = \int_0^{2\pi}\int_{-\pi/2}^{\pi/2} i_0(r,\theta,\phi)Y_{lm}^*(\theta,\phi)\sin\theta d\theta d\phi. \quad \text{(S16)}$$

##### 4.2.2 Time dependent fluorescence profile

For a perfect half bleach experiment, the concentration post-bleach is uniform ($i_0(r,\theta,\phi) = 1$) in the upper half of the compartment ($\theta > 0$) and 0 in the lower half ($\theta < 0$). As a results, the spherical harmonics amplitudes do no depend on $r$:

$$i_{lm,0} = \int_0^{2\pi} d\phi \int_0^{\pi/2} d\theta \; Y_{lm}^*(\theta,\phi)\sin\theta. \quad \text{(S17)}$$

In addition, because of the rotational symmetry of the post-bleach state, all modes with $m > 0$ have a zero amplitude. And because of the anti-symmetry with respect to the $\theta = 0$ plane, all modes where $l > 0$ and $l$ is even are also zero. So we only need to calculate:

$$B_{l0n,0} = i_{l0,0}\frac{\int_0^R j_l(k_{ln}r)r^2dr}{\int_0^R j_l^2(k_{ln}r)r^2dr}. \quad \text{(S18)}$$

In the end the time-dependent intensity profile after a half-compartment experiment is given by:

$$i(r,\theta,\phi,t) = B_{000,0} + \sum_{l=1,3,5...}^{\infty}\sum_{n=1}^{\infty} B_{l0n,0} Y_{l0}(\theta,\phi) j_l(k_{ln} r) e^{-Dk_{ln}^2 t}. \tag{S19}$$

The intensity profile is a sum of exponentially decaying modes. The dominant mode is the one corresponding to $l = 1$ and $n = 1$, for which $k_{11}R \simeq 2.08158$, meaning that $k_{11} \simeq 2.08158/R = 4.16316/L$. The characteristic decay time for this mode is thus $\tau_{11} \simeq 1/(Dk_{11}^2) = 0.0577L^2/D$. The normalized decay time is $\alpha_{L,\mathrm{sphere}} \simeq 0.577$.

### 5 Influence of frame acquisition interval on the precision and accuracy of the recovery time measurement

Additional simulations were performed to assess the respective influence of bleach duration and frame acquisition interval in the precision and accuracy of the data (Fig. S5).

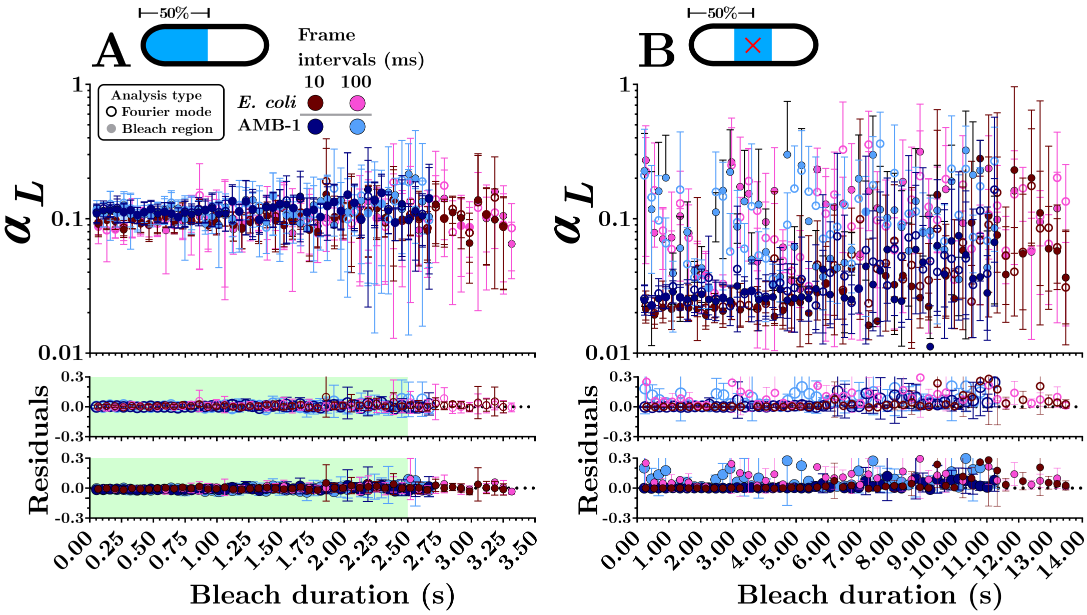


Figure S5: **Dependence of recovery time on photobleaching step duration.** Value of the coefficient $\alpha_L$ as a function of photobleaching step duration for (A) half-compartment and (B) central region (size of the central photobleached region $R = 1\,\mu m$) with a 10 ms and 100 ms frame interval. Simulated FRAP experiments were performed for cells with $L_T = 5\,\mu m$ and $D = 15\,\mu m^2\,s^{-1}$. In both (A) and (B), results are shown for cells with the shape and dimensions of a typical *E. coli* cell (shades of red) and of an *P. magneticum* cell (shades of blue), for the two different types of analysis considered. N = 500 per bacterium per analysis type.

#### 6 Influence of cell length on the value and precision of the recovery time extracted from experiments

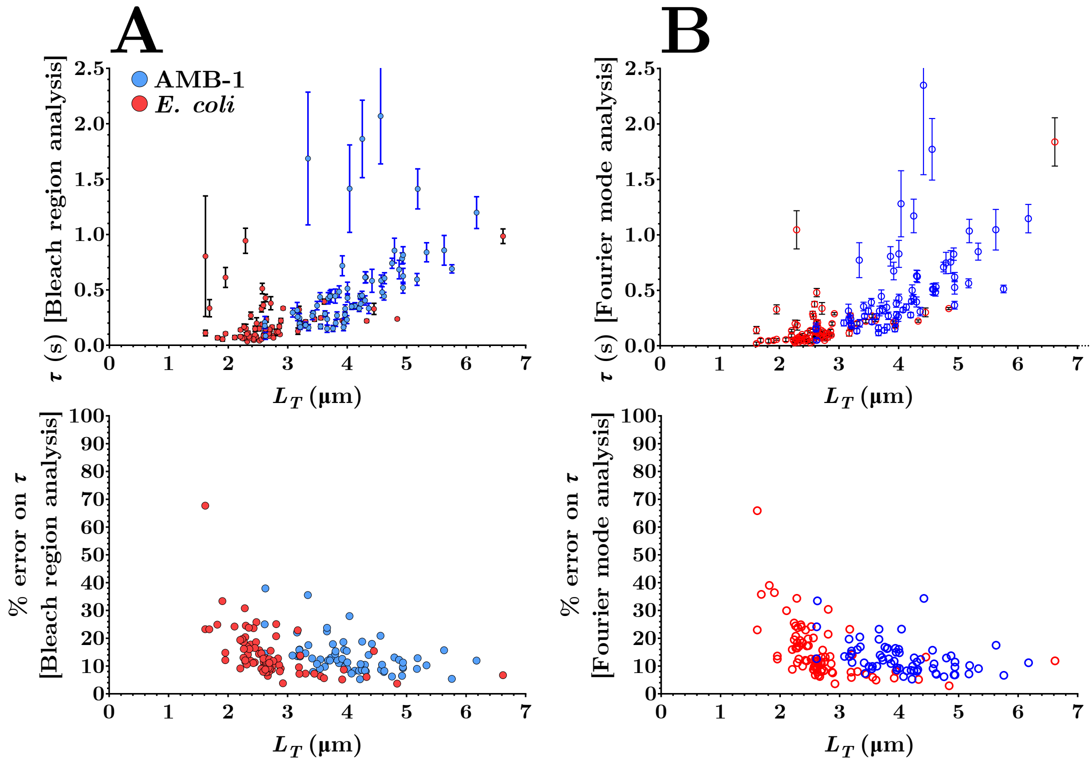


Figure S6: **Value and precision of experimentally measured recovery times** Recovery time $\tau$ (top panels) and percent error on the value of this recovery time obtained from the single-exponential fit of the experimental recovery curves for both *E.coli* (red symbols) and *P. magneticum* (blue symbols), using either (A) bleach region analysis or (B) Fourier mode analysis, plotted as a function of $L_T$. The error bars shown in the top panels for the value of $\tau$ are one standard deviation of error from the respective fits.

### References

[1] Lucas Le Nagard, Viviana Morillo-López, Cecile Fradin, and Dennis A Bazylinski. Growing magnetotactic bacteria of the genus Magnetospirillum: Strains MSR-1, AMB-1 and MS-1. *Journal of visualized experiments: JoVE*, (140):58536, 2018.

[2] Maiko Furubayashi, Andrea K Wallace, Lina M González, Justin P Jahnke, Brendan M Hanrahan, Alexis L Payne, Dimitra N Stratis-Cullum, Matthew T Gray, Han Liu, Melissa K Rhoads, et al. Genetic tuning of iron oxide nanoparticle size, shape, and surface properties in Magnetospirillum magneticum. *Advanced Functional Materials*, 31(4):2004813, 2021.

[3] Chen-Dong Yang, Haruko Takeyama, Tsuyoshi Tanaka, and Tadashi Matsunaga. Effects of growth medium composition, iron sources and atmospheric oxygen concentrations on production of luciferase-bacterial magnetic particle complex by a recombinant Magnetospirillum magneticum AMB-1. *Enzyme and Microbial Technology*, 29(1):13–19, 2001.
